# Factors that affect protein abundance of the bZIP transcription factor ABRE-BINDING FACTOR 2 (ABF2), a positive regulator of abscisic acid signaling

**DOI:** 10.1101/2021.01.25.428157

**Authors:** Katrina J Linden, Yi-Tze Chen, Khin Kyaw, Brandan Schultz, Judy Callis

**Author notes:** Correspondence: Judy Callis.

## Abstract

Most members of bZIP transcription factor (TF) subgroup A play important roles as positive effectors in abscisic acid (ABA) signaling during germination and/or in vegetative stress responses. In multiple plant species, one member, ABA INSENSITIVE 5 (ABI5), is a major transcription factor that promotes seed maturation and blocks early seeding growth in response to ABA. Other members, referred to as either ABRE-Binding Factors (ABFs), ABRE-Binding proteins (AREBs), or D3 PROTEIN BINDING FACTORS (DPBFs), are implicated as major players in stress responses during vegetative growth. Studies on the proteolytic regulation of ABI5, ABF1, and ABF3 in *Arabidopsis thaliana* have shown that the proteins have moderate degradation rates and accumulate in the presence of the proteasome inhibitor MG132. Exogenous ABA slows their degradation and the ubiquitin E3 ligase called KEEP ON GOING (KEG) is important for their degradation. However, there are some reported differences in degradation among subgroup A members. The conserved C-terminal sequences (referred to as the C4 region) enhance degradation of ABI5 but stabilize ABF1 and ABF3. To better understand the proteolytic regulation of the ABI5/ABFs and determine whether there are differences between vegetative ABFs and ABI5, we studied the degradation of an additional family member, ABF2, and compared its *in vitro* degradation to that of ABI5. As previously seen for ABI5, ABF1, and ABF3, epitope-tagged constitutively expressed ABF2 degrades in seedlings treated with cycloheximide and is stabilized following treatment with the proteasome inhibitor MG132. Tagged ABF2 protein accumulates when seedlings are treated with ABA but its mRNA levels do not increase, suggesting that the protein is stabilized in the presence of ABA. ABF2 is also an *in vitro* ubiquitination substrate of the E3 ligase KEG and recombinant ABF2 is stable in *keg* lysates. ABF2 with a C4 deletion degrades more quickly *in vitro* than full-length ABF2, as previously observed for ABF1 and ABF3, suggesting that the conserved C4 region contributes to its stability. In contrast to ABF2 and consistent with previously published work, ABI5 with C terminal deletions including an analogous C4 deletion is stabilized *in vitro* compared to full length ABI5. *In vivo* expression of an ABF1 C4 deletion protein appears to have reduced activity compared to equivalent levels of full length ABF1. Additional group A family members show similar proteolytic regulation by MG132 and ABA. Altogether, these results together with other work on ABI5 regulation suggest that the vegetative ABFs share proteolytic regulatory mechanisms that are not completely shared with ABI5.

## INTRODUCTION

The *Arabidopsis thaliana* family of basic leucine zipper (bZIP) transcription factors consists of 67-78 members (Jakoby et al. 2002, Deppmann et al. 2004, and Dröge-Laser et al. 2018). bZIP proteins are characterized by the presence of a DNA-binding basic region enriched in arginine or lysine residues and a leucine zipper dimerization motif that consists of a variable number of leucine heptad repeats. Arabidopsis bZIPs are divided into 13 groups (A-K, M, and S) defined by sequence similarity in the basic region and by shared conserved motifs outside of the bZIP domain (Dröge-Laser et al. 2018). Group A, with thirteen members, can be further divided into four subgroups (Dröge-Laser et al. 2018). Nine members in two subgroups share four additional regions called C(conserved)1-4 (Uno et al. 2000, Choi et al. 2000). The ABI5 subgroup consists of ABA INSENSITIVE 5 (ABI5, also referred to as Dc3 PROTEIN BINDING FACTOR or DPBF1) and three other DPBFs (DBPF2, DPBF3/AREB3 for ABA RESPONSIVE ELEMENT BINDING, and DPBF4/EEL for ENHANCED EM LEVEL). The next closest subgroup, with a diverged C4 region, is the ABF subgroup, consisting of ABF1, ABF2/AREB1, ABF3/DPBF5, ABF4/AREB2, and one member currently unnamed and uncharacterized (At5g42910) (Brocard et al. 2002, Dröge-Laser et al. 2018). With the exception of At5g42910, which has not been tested, the other eight members show binding to a synthetic multimer of an ABA-responsive element (Uno et al. 2000; Choi et al. 2000, Kim and Thomas 2002). Multiple members of group A are broadly represented among land plants and are also present in the bryophytes *Physcomitrella patens* and *Marchantia polymorpha*. Two *ABF/ABI5* members are identified in the charophyte *Klesormidium nitens*, which lacks an ABA-dependent response but does have a desiccation tolerance response, suggesting the origins of land plant ABA-dependent responses utilize group A bZIP proteins as an early component (Hauser et al. 2011, Cuming 2019).

*ABI5*, the best characterized group A member, was first identified from a mutant screen in Arabidopsis for ABA-insensitive germination (Finkelstein 1994, Finkelstein and Lynch 2000) and has emerged as a key player in ABA responses during germination and early seedling growth. *ABI5* transcripts increase during seed development with highest levels in dry seeds but are also present in vegetative tissues at much lower levels (Finkelstein and Lynch 2000, Lopez-Molina and Chua 2000, Brocard et al. 2002). There is a short window of time after seed stratification when exogenous ABA can strongly induce *ABI5* mRNA and protein accumulation (Lopez-Molina et al. 2001), which presumably inhibits germination and growth since *abi5* loss-of-function (LOF) mutants germinate in normally inhibitory concentrations of ABA (Finkelstein 1994; Lopez-Molina and Chua, 2000). Although at levels ~100-fold less than in 2-day old seedlings, ABA can modestly increase *ABI5* mRNA in 10-day old plants (Brocard et al. 2002). ABI5 does play roles during nitrate-mediated inhibition of lateral root growth (Signora et al. 2001) and in floral transition, the latter through positively modulating expression of *FLOWERING LOCUS C (FLC)* (Wang et al. 2013), a flowering repressing transcription factor (Michaels and Amasino 1999).

Members of the *ABF* subgroup appear to have more significant roles during vegetative stress responses. Multiple ABF mRNAs increase in response to exogenous salt and ABA in mature plants (Uno et al. 2000, Choi et al. 2000). *ABF1* (Yoshida et al. 2015), *ABF2*, *ABF3*, and *ABF4* (Yoshida et al. 2010) all contribute to drought tolerance in mature plants. Over-expression of ABF2 (Kim et al. 2004), ABF3 or ABF4 (Kang et al. 2002) slows germination, indicating that they can modulate this process as well.

In addition to regulation at the transcript level, ABI5/ABF abundance is regulated at the protein level (Liu and Stone 2014, Yu et al. 2015, Skubacz et al. 2016). Exogenous ABA slows the degradation of constitutively expressed ABI5 and ABF1 and ABF3 in transgenic seedlings (Lopez-Molina et al. 2001, Chen et al. 2013), and the RING-type E3 ligase KEEP ON GOING (KEG) plays an important role in ABI5, ABF1, and ABF3 degradation *in vivo* and ubiquitinates them *in vitro* (Stone et al. 2006, Chen et al. 2013). Additional ubiquitin ligases are implicated in promoting ABI5 degradation. DWA1 (DWD HYPERSENSITIVE TO ABA 1) and DWA2 are two substrate specificity factors for a Cullin4-based ubiquitin E3 ligase (Lee et al. 2010) whose LOF mutants have ABA-hypersensitive phenotypes. Both interact *in vitro* with ABI5 and *dwa* extracts degrade ABI5 more slowly. ABA-HYPERSENSITIVE DCAF 1 (ABD1), another CUL4 substrate receptor, interacts with ABI5 and plays a role in ABI5 degradation (Seo et al. 2014). ABD1 and ABI5 interact in a yeast two-hybrid assay, and endogenous ABI5 pulled down with Myc-ABD1 in a co-IP from lysate of plants overexpressing Myc-ABD1. When seeds were treated with CHX following ABA treatment, ABI5 protein degraded more slowly in *abd1-1* seeds than in wild-type seeds (Seo et al. 2014). With the exception of KEG, the role of these ligases in the proteolytic control of other group A bZIPs is not known.

The earliest described Arabidopsis ABI5/ABF post-translational modification is phosphorylation; using in-gel kinase assays, a 42-kDa kinase activity modifying peptide substrates corresponding to ABF2 or ABF4 C1, C2, or C3 regions is rapidly activated after *in vivo* ABA treatment (Uno et al. 2000, Furihata et al. 2006). Arabidopsis ABF *in vivo* phosphorylation at two sites increased after treatment of leaves with ABA; one corresponds to an AREB3 C3 serine, but the other phosphorylated C2-Ser cannot be assigned to a specific ABF because the tryptic peptide containing the site is 100% conserved among 7 ABF members (Kline et al. 2010). Interestingly, no net change in constitutively expressed Arabidopsis ABI5 phosphorylation is observed after ABA treatment of 8-day-old seedings, although migration on SDS-PAGE is affected (Lopez-Molina et al. 2001, Lopez-Molina et al. 2002). Peptides that include the same C1-3 regions, but not C4, were identified as singly phosphorylated in HA-ABI5 by mass spectrometry from 4-week-old plants expressing ABI5 under the 35S promoter (Lopez-Molina et al. 2002). SnRK2 kinases at ~42-44 kDa were identified as ABA-inducible activities in a number of species (Johnson et al. 2002, Nakashima et al. 2009). At least two other kinases were observed in in-gel kinase assays (Johnson et al. 2002, Nakashima et al. 2009), leading to identification of calcium-dependent protein kinases (CDPKs), named CPKs in Arabidopsis. CPK32 interacts with ABFs 1-4 and phosphorylates an ABF4 peptide *in vitro* (Choi et al. 2005). CPK10 and 30 interact in yeast with ABF4 (Choi et al. 2005) and CPK4 and 11 phosphorylate full-length ABF1 and ABF4 in in-gel assays (Zhu et al. 2007). A calcinurin-type kinase, CIPK26, interacts with ABI5 and phosphorylates it *in vitro* (Lyzenga et al. 2013). Recently, another CDPK, CPK6, was shown to interact with ABFs1-4 and ABI5 *in planta* and phosphorylation *in vitro* of ABI5 and ABF3 was demonstrated (Zhang et al. 2020). These studies have revealed a complex network of phosphorylations, with additional kinases likely awaiting discovery and characterization. A phospho-mimic form of AREB1/ABF2 with five Ser to Asp substitutions trans-activates ABRE-containing genes in an ABA-independent manner, suggesting that ABA-dependent phosphorylation activates the transcriptional activation function (Furihata et al. 2006). Recombinant ABFs, which are presumed to be unphosphorylated, bind to ABREs *in vitro* (Uno et al. 2000), in yeast one-hybrid assays (Choi et al. 2000) and are active in yeast (Kim et al. 2002), so the precise activating mechanism remains unclear. Current evidence suggests that phosphorylation status does not affect protein longevity: ABI5 with C1-4 serine/threonine residues substituted either for alanine or aspartate were equivalently degraded *in vitro* (Liu and Stone 2014).

Other post-translational modifications of ABFs/ABI5 have been linked to protein longevity. ABI5 is S-nitrosylated in an NO-dependent manner, and this modification negatively affects ABI5 accumulation (Albertos et al. 2015). The SUMO-modification pathway, another protein modification system, also modulates ABA responses. A reduction in sumoylation components increases ABA sensitivity in seed germination and root growth, and ABI5 mono-sumoylation *in vivo* is dependent on SIZ1, the major plant SUMO E3 ligase (Miura et al. 2009). The genetic relationship between *siz1* and *abi5* mutants suggest that SIZ1 suppresses ABA responses in an ABI5-dependent manner.

There are some differences and unknowns in the proteolytic regulation of ABFs. Of the seven group A members, only DPBF2 retains the cysteine required for S-nitrosylation, and whether ABFs are sumoylated is relatively uncharacterized. ABFs lack the SUMO consensus sequence found in ABI5 (using sumoplot) and some members lack a similarly located lysine SUMO attachment site identified in ABI5. However, ABF3 was positive in a global screen for SUMO targets enriched after heat stress, although the attachment site was not reported (Rytz et al. 2018). Another example of a difference in proteolytic regulation among group A proteins is the role of the C4 region. These sequences enhance degradation of ABI5 (Liu and Stone 2013) but slow ABF1 and ABF3 degradation in *in vitro* assays (Chen et al. 2013). ABA-dependent phosphorylation at a conserved serine/threonine in the C4 region has been proposed to play a role in stabilizing ABF3 by promoting ABF3 interaction with a 14-3-3 protein (Sirichandra et al. 2010). This phosphorylation site is conserved in ABI5 and also plays a role in ABI5 interaction with 14-3-3 proteins. In barley, HvABI5 interacts with several 14-3-3 proteins, and introducing a phospho-null substitution at T350 in the C4 region in HvABI5 eliminated interaction with 14-3-3 proteins in a yeast two-hybrid assay (Schoonheim et al. 2007). Arabidopsis ABI5 also interacts with a 14-3-3 protein in a yeast two-hybrid assay (Jaspert et al. 2011). While the C4 phosphorylation site appears to play similar roles in 14-3-3 binding for ABI5 and ABF3, sequence conservation between ABI5 and the ABFs is low in this region and could account for the different role of the C4 in regulating ABI5 compared to the other ABFs.

To better characterize ABF proteolytic regulation and to understand the proteolytic differences between ABF and ABI5 subgroups, we characterized the degradation of an additional ABF family member, ABF2. Like ABF1 and ABF3, ABF2 degrades in seedlings treated with cycloheximide, is stabilized following treatment with the proteasome inhibitor MG132, rapidly accumulates in seedlings treated with ABA, and is an *in vitro* KEG substrate. ABF2 with a C4 deletion degrades more quickly than full-length ABF2 *in vitro*, while ABI5 degradation in the same assays is slowed by C-terminal deletions, showing that the C4 region of ABF2 contributes to its stability and confirming differences between ABI5 and ABFs. We also characterized the *in vivo* activity of a C4 deletion form of ABF1. Seeds from lines overexpressing full-length ABF1 germinated more slowly than seeds from lines overexpressing ABF1 with a C4 deletion, suggesting that the C4 region is important for ABF1 activity. A survey of other ABF/ABI5 subgroup members suggests that their regulation by ABA and the proteasome is similar to that of previously characterized ABFs. Altogether, these results suggest that the members of the ABF share proteolytic regulatory mechanisms that are not completely shared with ABI5.

## MATERIALS AND METHODS

### Plant growth conditions, materials, and genotyping

*A. thaliana* seeds were surface-sterilized in a solution of 25% bleach and 0.02% Triton-X100 for ten minutes, rinsed with sterile H_2_O, and stratified at 4°C for at least 24 hours before plating. For agar-grown seedlings, seeds were plated on solid growth media (GM) consisting of g/L Murashige and Skoog basal salt mixture (Sigma), 1% sucrose, 0.5 g/L MES, 1X B vitamins (0.5 μg/mL nicotinic acid, 1 ug/mL thiamine, 0.5 μg/mL pyridoxine, and 0.1 μg/mL myo-inositol, Sigma), and 8 g/L Bacto Agar (Becton Dickinson), pH 5.7. After two weeks at room temperature under constant light, seedlings were transplanted from agar GM plates to soil, and plants were grown at 20°C with 50% humidity and 16 hours light/8 hours dark. For liquid-grown seedlings, approximately 100 seeds were added to 1 mL liquid GM distributed around the periphery of a 60 mm petri dish (Falcon #353002) and grown at room temperature under constant light.

*A. thaliana* ecotype Col-0 (CS70000) and *keg* insertion alleles were obtained from the Arabidopsis Biological Resource Center in Columbus, Ohio (https://abrc.osu.edu/). The *keg-1* (At5g13530) T-DNA line SALK_049542 and the *keg-2* T-DNA line SALK_018105 (previously characterized in Stone et al. 2006) were maintained as heterozygotes because homozygous individuals do not progress beyond the seedling stage. Homozygous *keg* seedlings were identified by their phenotypic differences from wild-type siblings (Stone et al. 2006). The *keg* genotype was verified by PCR using primers listed in Supplemental **Table 1**. For *keg-1* lines, primer 5-253 was used with primer 5-254 to produce a WT gene-specific product, and primer 5-254 was used with the T-DNA left border primer 9-001 for a T-DNA junction product. For *keg-2* lines, primer 9-113 was used with primer 9-114 for a WT gene-specific product, and primer 9-114 was used with the T-DNA left border primer 9-001 for a T-DNA junction product.

To generate transgenic *A. thaliana* lines expressing epitope-tagged proteins, the plant expression constructs described below were transformed into *Agrobacterium tumefaciens* strain AGL1. The floral dip method (Clough and Bent 1998) was used to transform the expression construct into Col-0 plants. Transformants that segregated 3:1 on GM with 50 μg/mL kanamycin were propagated and homozygous T3 or T4 seedlings were used in experiments (except for Figures 2.1, 2.2 and Supplemental Figures 2-4, where T2 segregating seed was used). Transgenic line information is in Supplemental Table 4.

### Cycloheximide, MG132, and ABA treatments

Seedlings were grown in sterile liquid GM in 60 mm petri dishes and after six days the liquid GM was replaced to allow seedlings to adjust to fresh media overnight. The following day, for cycloheximide treatments the liquid GM was replaced with liquid GM containing 0.2 mg/mL cycloheximide (Sigma) or plain liquid GM as a mock control. For MG132 treatments, the liquid GM was replaced with liquid GM containing 50 μM MG132 (Peptides International #IZL-3175-v, or Selleck Chemical #S2619-5MG) or 0.5% DMSO as a solvent control. For ABA treatments, the liquid GM was replaced with liquid GM containing 50 μM ABA (Sigma) or 0.1% EtOH as a solvent control. After the indicated time, seedlings were flash frozen in liquid nitrogen and ground in fresh buffer (50 mM Tris pH 8.1, 250 mM NaCl, 0.5% NP-40, 1 mM PMSF, 50 μM MG132, and 1 cOmplete Mini EDTA-free protease inhibitor tablet (Roche #11836170001) per 10 mL buffer). Lysate was collected after centrifugation at 13,000 rpm for 10 minutes at 4°C, and total protein concentration was measured using a Bradford assay (Protein Assay Dye Reagent Concentrate, Bio-Rad). Equal amounts of total protein per sample were separated by 10% SDS-PAGE, and 10xMyc-ABF2 protein was visualized with Anti-myc western blotting. Anti-actin western blotting or Ponceau S staining of the membrane (Supplemental Figures 2-4) was used to visualize actin for a loading control.

### Cell-free degradation assays

Cell-free degradation assays were performed based on those described in Wang et al. 2009. Approximately 100 7-day-old liquid grown seedlings (0.15g fresh weight) were flash frozen in liquid nitrogen and stored at −80°C until use. Each 0.15g tube of frozen seedlings was ground in 350 μL fresh buffer (25 mM Tris, 10 mM ATP, 10 mM MgCl_2_, 10 mM NaCl, and 1 mM DTT). Lysate (approximately 2.5 μg/μL total protein) was collected after centrifugation at 13,000 rpm for 10 minutes at 4°C. 300 μL lysate was incubated at 22°C in a thermocycler and approximately 2 μg of recombinant protein purified from *E. coli* was added to the lysate. For assays comparing degradation of multiple proteins, one lysate was divided into 300 uL aliquots. For assays with MG132, 1.5 μL of 10 mM MG132 in DMSO, or 1.5 μL of DMSO as a solvent control, was added to the lysate prior to the addition of recombinant protein. Samples were removed at indicated time points, mixed with Laemmli sample buffer, and heated to 98°C for 7 minutes. Proteins were separated with 10% SDS-PAGE and anti-FLAG, anti-HA, anti-Myc, or anti-GST Western blotting was used to visualize the recombinant protein in each sample.

### Germination experiments

Age matched seeds were sterilized and plated on GM as described above, stratified for three days at 4°C, then incubated at 20°C in the light. At specific time points, the plates were scored for germination (radicle emergence) and returned to the growth chamber. Time courses for each line were repeated 3 times with ~50 seeds per experiment.

### RNA extraction and quantitative PCR (qPCR)

Seedlings were grown in liquid GM under constant light for 7 days and treated for six hours with 50 μM ABA or 0.1% ethanol as a solvent control. Total RNA was isolated using the RNeasy Plant Mini Kit (Qiagen, 74904) according to manufacturer’s instructions. 2 μg of total RNA was used in each 10 μl reverse transcription reaction performed with the SuperScript III First-Strand Synthesis System for RT PCR (Invitrogen, 18080-051) according to manufacturer’s instructions. Real-time PCR amplification was performed with 20 μl reactions containing 2 μl of first-strand cDNA (diluted 1/10 with H_2_O), 10 pmoles of each primer, and 10 μl PowerUp SYBR Green Master Mix (Applied Biosystems, A25742). Relative transcript levels were obtained using the comparative Ct method, normalized to *ACT2*. The experiment was performed independently 3 times with 3 technical replicates each time. Primer sequences are listed in **Supplemental Table 5**.

### Cloning and constructs

Primer sequences for plant expression cloning are listed in **Supplemental Table 2**. For the plant expression construct for 10xMyc-ABF2 under control of the *35S* promoter (p9186), plasmid pVM491 (*ABF2* cDNA in pENTR223, stock # G85579) was obtained from the Arabidopsis Biological Resource Center (ABRC) in Columbus, Ohio (https://abrc.osu.edu/). A Gateway (Invitrogen) LR reaction was used to recombine the *ABF2* CDS into pGWB21 (pVM259) (Nakagawa et al. 2007) following manufacturer protocols. Similarly, cDNAs for *DPBF2* (G20103, pVM493) and *DPBF4* (G83893, pVM492) were obtained from the ABRC and recombined into pGWB21. For *ABF4* and *AREB3*, RNA was isolated from Arabidopsis seedlings and cDNAs were synthesized as described above, amplified with primers 9-363 and 9-364 (*ABF4*) or 9-365 and 9-366 (*AREB3*), cloned into pDONR201 with Gateway BP reactions, sequence verified, and recombined into pGWB21 with Gateway LR reactions. For expression of His_6_-HA_3_-ABF1 (p9204) and His_6_-HA_3_-ABF1-ΔC4 (p9205) under control of the *UBQ10* promoter, cDNAs synthesized as described above were amplified with primers 9-438 and 9-439 (*ABF1*) or 9-438 and 9-452 (*ABF1-ΔC4).* The PCR products were digested with AseI and BamHI, then ligated into p3756, a modified pGreenII plasmid (Gilkerson et al. 2015), that had been cut with NdeI and BamHI. The line expressing GFP under control of the *UBQ10* promoter (line 13240) was described in (Dreher 2006).

Primer sequences for bacterial expression cloning are listed in **Supplemental Table 3**. For the bacterial expression construct for full-length His_6_-FLAG-ABF2 (p9193), a Gateway LR reaction was used to recombine the *ABF2* cDNA from G85579 into the bacterial expression vector pEAK2 (Kraft 2007).

For the bacterial expression construct for full-length His_6_-HA_3_-ABF2 (p9213), p9208 (*ABF2* cDNA cloned into a modified pGreenII plasmid using primers 9-440 and 9-441) was digested with *NdeI* and *BamHI*. The *ABF2* sequence was ligated into NdeI and BamHI sites in the bacterial expression vector p3832, a modified pET3c (Novagen) plasmid containing a 5’ His_6_-HA_3_ cassette.

For the bacterial expression construct for His_6_-HA_3_-ABF2-ΔC4 (p6881), p9193 (*ABF2* cDNA in pEAK2, described above) was used as a template and primers 9-440 and 6-1072 were used to amplify the *ABF2* sequence, introduce a stop codon after base 1245 (relative to A of translation start ATG), and add *NdeI* and *BamHI* restriction sites to the 5’ and 3’ ends, respectively. The sequence was ligated into *NdeI* and *BamHI* sites in the bacterial expression vector p3832.

For the bacterial expression constructs for His_6_-HA_3_-ABI5 (p6943), -ABI5^1-343^ (p6944), and -ABI5-ΔC4 (p6945), *ABI5* cDNA in pDONR (plasmid p9017, derived from the ABRC clone PYAt2g36270 described in Stone et al. 2006) was used as a template and primer 6-1107 was used with 6-1109 (ABI5), 6-1108 (ABI5^1-343^), or 6-1110 (ABI5-ΔC4) to amplify the *ABI5* sequence, introduce stop codons after base 1029 for ABI5^1-343^ or base 1287 for -ΔC4, and add *NdeI* and *BamHI* restriction sites to the 5’ and 3’ ends, respectively. The *ABI5*, *ABI5^1-343^*, and *ABI5-ΔC4* sequences were ligated into *Nde* I and *Bam* HI sites in the bacterial expression vector p3832.

The Myc-TGA1 (p9109) and Myc-GBF3 (p9107) constructs contain *TGA1* or *GBF3* cDNAs in p7296, a pEXP1-DEST expression vector (Thermo) modified with a Myc9 tag. RNA was isolated from Arabidopsis seedlings and cDNAs were synthesized as described above, amplified with primers 9-367 and 9-368 (*TGA1*) or 9-361 and 9-362 (*GBF3*), recombined into pDONR with Gateway BP reactions, then recombined with Gateway LR reactions into p7296. Sequences were verified. The GST-EGL3-FLAG construct (pVM567) contains *EGL3* in the pGEX-4T-1 vector. It was a gift from Ling Yuan at the University of Kentucky. The GST-FUS3 construct (pVM505) was obtained from Sonia Gazarrini as described in Chen et al. 2013. The GST-KEG-RKA construct was described in Stone et al. 2006. The UBC10 E2 construct was described in Kraft et al. 2005.

### Recombinant protein expression

*E. coli* transformed with expression constructs was grown in 10 mL cultures overnight at 37°C, then added to flasks containing 500 mL LB (10 g/L tryptone, 5 g/L yeast extract, 5 g/L NaCl, 15 g/L agar, pH 7.5). The 500 mL cultures were grown at 37°C for approximately 3 hours until the OD_600_ was around 0.4−0.6, at which point the cultures were moved to room temperature and protein expression was induced with the addition of 0.5 mM isopropyl β-D-1-thiogalactopyranoside (IPTG) for three to four hours. Cells were collected by centrifugation and pellets were stored at −80°C. *E. coli* strains and induction compounds were as follows. His_6_-HA_3_-ABF2-ΔC4, His_6_-HA_3_-ABI5, -ABI5^1-343^, and -ABI5-ΔC4, Myc-GBF3, Myc-TGA1, and GST-EGL3-FLAG were expressed in BL21(DE3)pLysS cells and expression was induced with IPTG. His_6_-HA_3_-ABF2 and His_6_-FLAG-ABF2 were expressed in Lemo21(DE3) cells and induced with IPTG. GST-FUS3 was expressed as described in (Chen et al. 2013).

### Protein purification/enrichment

Cell pellets were thawed on ice and resuspended in lysis buffer (25 mM Tris pH 7.5, 500 mM NaCl, 0.01% Triton-X100, 30 mM imidazole, and 1 cOmplete Mini EDTA-free protease inhibitor tablet per 50 mL buffer). Cells were lysed by sonication and supernatant was reserved after centrifugation. His- or GST-tagged proteins were captured from the lysate by the addition of 100 μL Ni Sepharose High Performance beads (GE Healthcare) or Glutathione Sepharose High Performance beads (GE Healthcare). Myc-tagged proteins were captured by addition of 75 μL EZview Red Anti-c-Myc Affinity Gel beads (Sigma). Beads were collected with centrifugation and rinsed at least 3 times with wash buffer (25 mM Tris pH 7.5, 300 mM NaCl, and 0.01% Triton-X100). Proteins were eluted from the beads by addition of 300 μL elution buffer (25 mM Tris, 150 mM NaCl, 0.01% Triton-X100, and 300 mM imidazole for His-tagged proteins or 20 mM reduced glutathione for GST-tagged proteins) and shaking at 4°C for 30 minutes. Myc-tagged proteins were eluted with 100 ug/mL c-Myc peptide in 250 μL RIPA buffer. Supernatant was collected after centrifugation and mixed with glycerol for a final concentration of 20% glycerol, and frozen at −80°C.

### In vitro ubiquitination assays

Substrate was incubated with E1, E2, E3, and ubiquitin in buffer (50 mM Tris-HCl pH 7.5, 10 mM MgCl_2_, 1 mM ATP, 200 μM DTT, and 2.1 mg/mL phosphocreatine) for 2 hours at 30°C. Each 30 μL reaction included approximately 50 ng of yeast E1 (Boston Biochem), 250 ng of E2 (AtUBC10), 5-10 μL of bead-bound E3 or 2-5 μL of soluble E3 (GST-KEG-RKA), 4 μg ubiquitin (Sigma), and 2 μL substrate (His_6_-HA_3_-ABF2). To stop the reaction, 10 μL of 5X LSB was added and samples were boiled for 10 minutes.

### SDS-PAGE and Western blotting

Proteins were separated on 10% polyacrylamide gels and transferred onto PVDF-P membranes (Immobilon). Membranes were blocked in 5% nonfat powdered milk (Carnation) dissolved in TBS + 0.1% Tween 20 (Sigma). Proteins were detected with the following antibodies. FLAG-tagged proteins were detected with Monoclonal ANTI-FLAG M2-Peroxidase (Sigma #A8592) at a dilution of 1:5,000. HA-tagged proteins were detected with Anti-HA-Peroxidase (Roche #12013819001) at a dilution of 1:5,000. Myc-tagged proteins were detected with Anti-c-myc-Peroxidase (Roche #11814150001) at dilution of 1:5,000. GST-tagged proteins were detected with rabbit polyclonal GST (Z-5): sc-459 (Santa Cruz Biotechnology) at a dilution of 1:5,000, followed by the secondary antibody peroxidase-conjugated AffiniPure Goat Anti-Rabbit IgG (H + L) (Jackson ImmunoResearch) at a dilution of 1:10,000. Actin was detected with Monoclonal Anti-Actin (plant) antibody produced in mouse (Sigma #A0480-200UL) at a dilution of 1:5000, followed by peroxidase-labeled Antibody to Mouse IgG (H + L) produced in goat (KPL #074-1806) at a dilution of 1:10,000. SuperSignal West Pico Chemiluminescent Substrate (Thermo Scientific #34078) or ProSignal Dura (Prometheus #20-301) were used as chemiluminescent substrates. Chemiluminescence was captured on X-ray film (Phenix) or digitally imaged in the linear range of detection using the ImageQuant LAS400 imaging system (GE). For quantitation of *in vivo* degradation or accumulation, ABF values were normalized to actin and treatment values were divided by control values. Curve fitting, half-life determinations, and t-tests were performed using Prism (GraphPad) as described in each figure legend. In GraphPad, one-phase decay is the term for first order decay.

## RESULTS

### ABF2 degrades in seedlings and increases following MG132 treatment

To examine the proteolytic regulation of ABF2 in plants, independent transgenic lines expressing Myc-ABF2 under control of the *35S* promoter were generated. 7-day-old seedlings from three independent lines were treated with the protein synthesis inhibitor cycloheximide (CHX). Once translation is inhibited by CHX, no new ABF2 protein is synthesized in the seedlings and degradation of the existing ABF2 protein can be visualized by monitoring protein levels over time. In all three lines, Myc-ABF2 protein was reduced over time following CHX treatment, indicating that Myc-ABF2 is unstable *in vivo* (**Figure 1A and Supplemental Figure 1A-C**). Proteasomal substrates typically accumulate after proteasome activity is inhibited by the cell-permeable substrate analog MG132. Myc-ABF2 protein increased when intact seedlings were treated with MG132 (**Figure 1B and Supplemental Figure 1D**), indicating that Myc-ABF2 degradation requires the proteasome.

**Figure 1:**
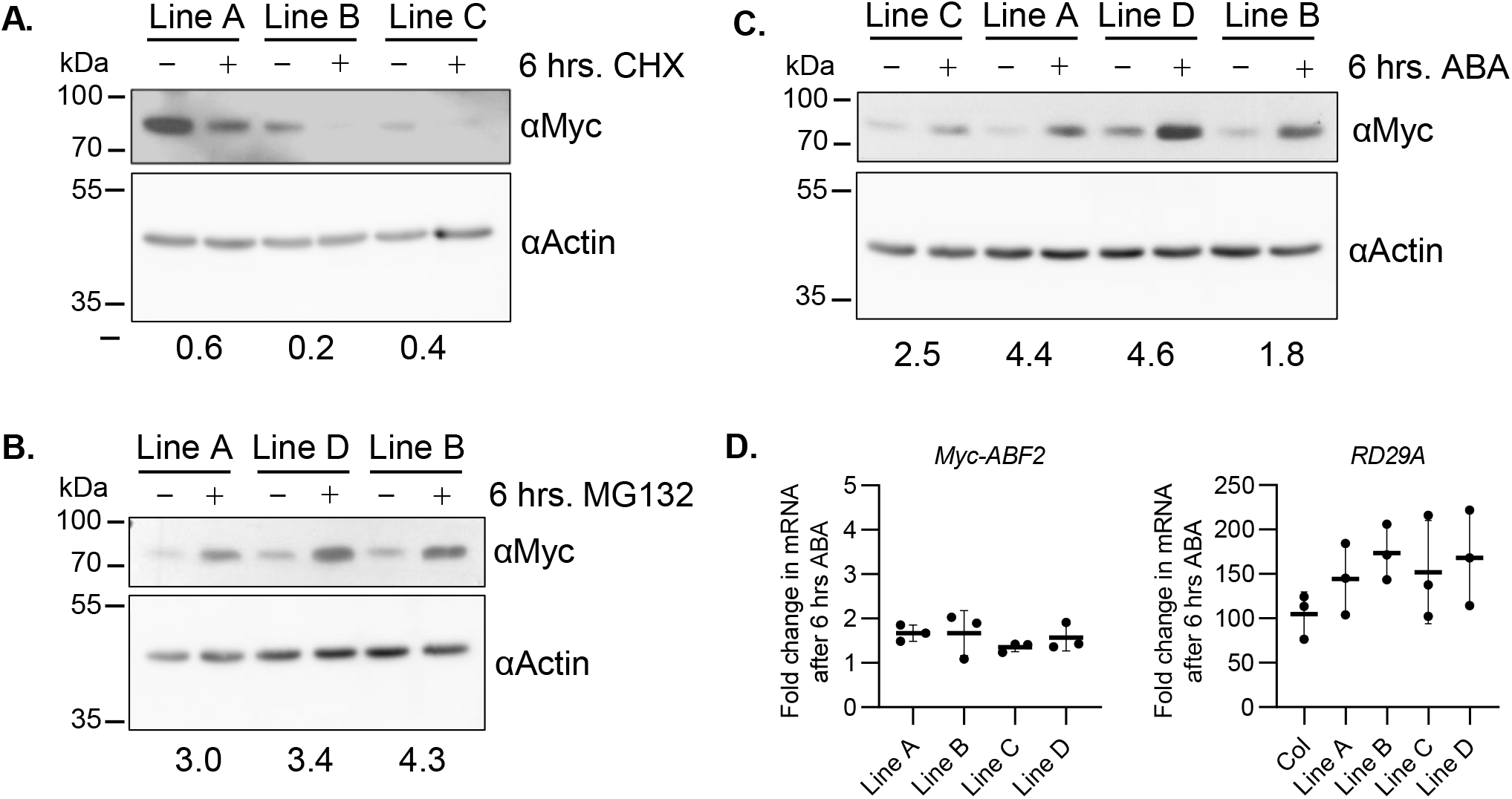
In seedlings, ABF2 protein decreases following cycloheximide treatment and increases following MG132 or ABA treatment. 7-day-old liquid grown seedlings from independent lines expressing 10xMyc-ABF2 were treated with **(A)** 0.2 mg/mL CHX dissolved in GM or plain liquid GM as a solvent control for six hours, **(B)** 50 μM MG132 in DMSO or 0.5% DMSO as a solvent control for six hours, or **(C)** with 50 μM ABA in EtOH or 5% EtOH as a solvent control for six hours. Equal amounts of total protein per sample were loaded for SDS-PAGE and anti-Myc western blotting was used to visualize the 10xMyc-ABF2 protein in each sample (**upper panels**). For a loading control, anti-actin western blotting was performed on the same membranes (**lower panels**). To measure the change in ABF2 protein for each line, all bands were quantified in Image Studio Lite, ABF2 values were normalized to actin values, and treatment value was divided by control. The treatment/control ratio is reported below each pair of samples. **D)** Expression of *Myc-ABF2* and *RD29A* mRNAs in 7-day-old seedlings treated with ABA for six hours, relative to mRNAs in mock-treated seedlings. qPCR results are shown as mean ± SD. N = three biological replicates with 3 technical replicates each. Data were normalized to *ACT2. RD29A* serves as a positive control for ABA treatment. Note: the seedlings used in A are from the homozygous T3 generation, the seedlings used in B and C are from the segregating T2 generation, and the seedlings used in D are from the homozygous T4 generation.

### ABF2 increases in seedlings following ABA treatment

Studies have shown that levels of constitutively expressed ABI5, ABF1, and ABF3 proteins increase in plants in response to exogenously supplied ABA (Lopez-Molina et al. 2001, Chen et al. 2013). To test whether ABF2 behaves similarly, seedlings from four independent transgenic lines expressing Myc-ABF2 under control of the *35S* promoter were treated with ABA (in both the segregating T2 and homozygous T4 generations). Myc-ABF2 protein increased in all four lines following ABA treatment (**Figure 1C and Supplemental Figure 1E,F**). To support the hypothesis that this accumulation results from protein stabilization and not an increase in synthesis, the mRNA levels of the *Myc-ABF2* transgene were measured after 6 hrs of ABA or after six hours of 0.1% EtOH as a solvent control. While mRNA for the positive control *RD29A* (Yamaguchi-Shinozaki and Shinozaki 1993) increased more than 100-fold after ABA treatment, *Myc-ABF2* mRNAs did not increase as much as Myc-ABF2 protein levels (Figure 1D).

### Survey of other group A bZIP proteins reveals regulation by MG132 and ABA

To investigate the proteolytic stability of other group A members, analogous epitope-tagged ABF4, DPBF2, DPBF4, and AREB3 proteins were expressed from the same vector system described above, and three independent transgenic lines overexpressing each protein were tested for effects of MG132 and ABA on protein accumulation (Supplemental figures 2, 3, and 4). ABF4, DPBF2, and DPBF4 accumulated in the presence of MG132 (AREB3 lines were not tested), and all four (ABF4, DPBF2, DPBF4, and AREB3) accumulated in the presence of ABA.

### *ABF2 degrades rapidly* in vitro *and is stabilized by MG132*

To test whether ABF2 protein is unstable *in vitro*, we used a cell-free degradation assay (as described in Wang et al. 2009) in which recombinant ABF2 protein expressed in and purified from *E. coli* was added to a lysate from wild-type Arabidopsis seedlings. Samples were removed from the reaction at various timepoints, and Western blotting was used to visualize the amount of tagged ABF2 protein remaining at each timepoint. We tested ABF2 with two different epitope tags (HA and FLAG) to show that ABF2 protein degradation *in vitro* does not depend on the nature of the tag. Both His_6_-HA_3_-ABF2 and His_6_-FLAG-ABF2 degraded quickly in the cell-free degradation assay and were stabilized by the addition of MG132 (**Figure 2**). One-phase decay curves predicted similar half-lives of 2.3 minutes and 2.8 minutes for His_6_-HA_3_-ABF2 and His_6_-FLAG-ABF2, respectively (**Figure 2 C, D**).

**Figure 2:**
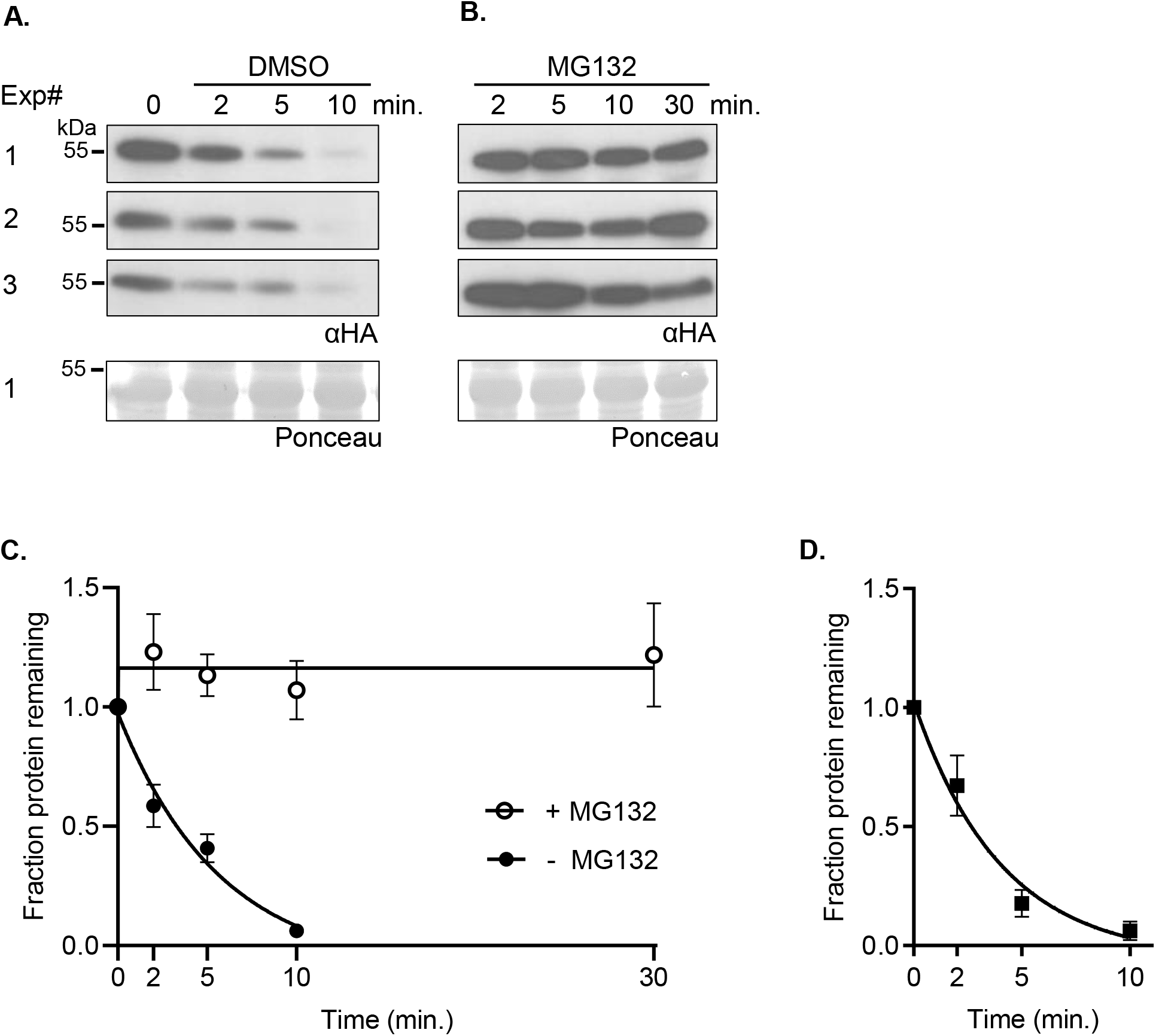
ABF2 degrades *in vitro* and is stabilized by MG132. In a cell-free degradation assay, recombinant full-length **(A-C)** His_6_-HA_3_-ABF2 or **(D)** His_6_-FLAG-ABF2-FL was added to lysate from Col seedlings and the reaction was split into separate tubes with equal volumes of either **(A)** DMSO or **(B)** 200 μM MG132 dissolved in DMSO and incubated at 22°C. Samples were removed at the indicated times, mixed with Laemmli sample buffer, and heated to 98°C to stop the reaction. Three independent experiments were performed. **(A, B)** SDS-PAGE followed by anti-HA or anti-FLAG western blotting was used to visualize the ABF2 protein in each sample (upper panels). Ponceau staining was used to demonstrate equal loading (lower panels). Ponceaus for Experiment #1 are shown. **(C)** Western blots were quantified with Image Studio Lite and the fraction of protein remaining at each timepoint was graphed. For each reaction, GraphPad Prism nonlinear regression was used to fit a curve modeling one-phase decay of ABF2. R squared is 0.93. Bars are SE. N = 3 independent experiments. **(D)** Graph of identical time course with recombinant His_6_-FLAG-ABF2-FL (western blots not shown). R squared is 0.91. Bars are SE. N = 3 independent experiments.

### *ABF2 is ubiquitinated by KEG* in vitro, *and KEG affects ABF2 degradation*

The E3 ligase KEG ubiquitinates ABI5, ABF1, and ABF3 *in vitro*, and promotes their degradation *in vivo* (Stone et al. 2006, Liu and Stone 2013, Chen et al. 2013). Degradation of recombinant ABF1 and ABF3 is slowed in *keg* lysates (Chen et al. 2013), and endogenous ABI5 hyperaccumulates in *keg* seedlings (Stone et al. 2006). We used an *in vitro* ubiquitination assay to test whether ABF2 is also a potential *in vivo* KEG substrate. When recombinant His_6_-HA_3_-ABF2 was incubated with a recombinant form of KEG and other proteins required for ubiquitination (E1, E2, and free ubiquitin), higher migrating forms of His_6_-HA_3_-ABF2 were observed only in the complete reaction (**Figure 3**), indicating covalent ubiquitin attachment to His_6_-HA_3_-ABF2.

**Figure 3:**
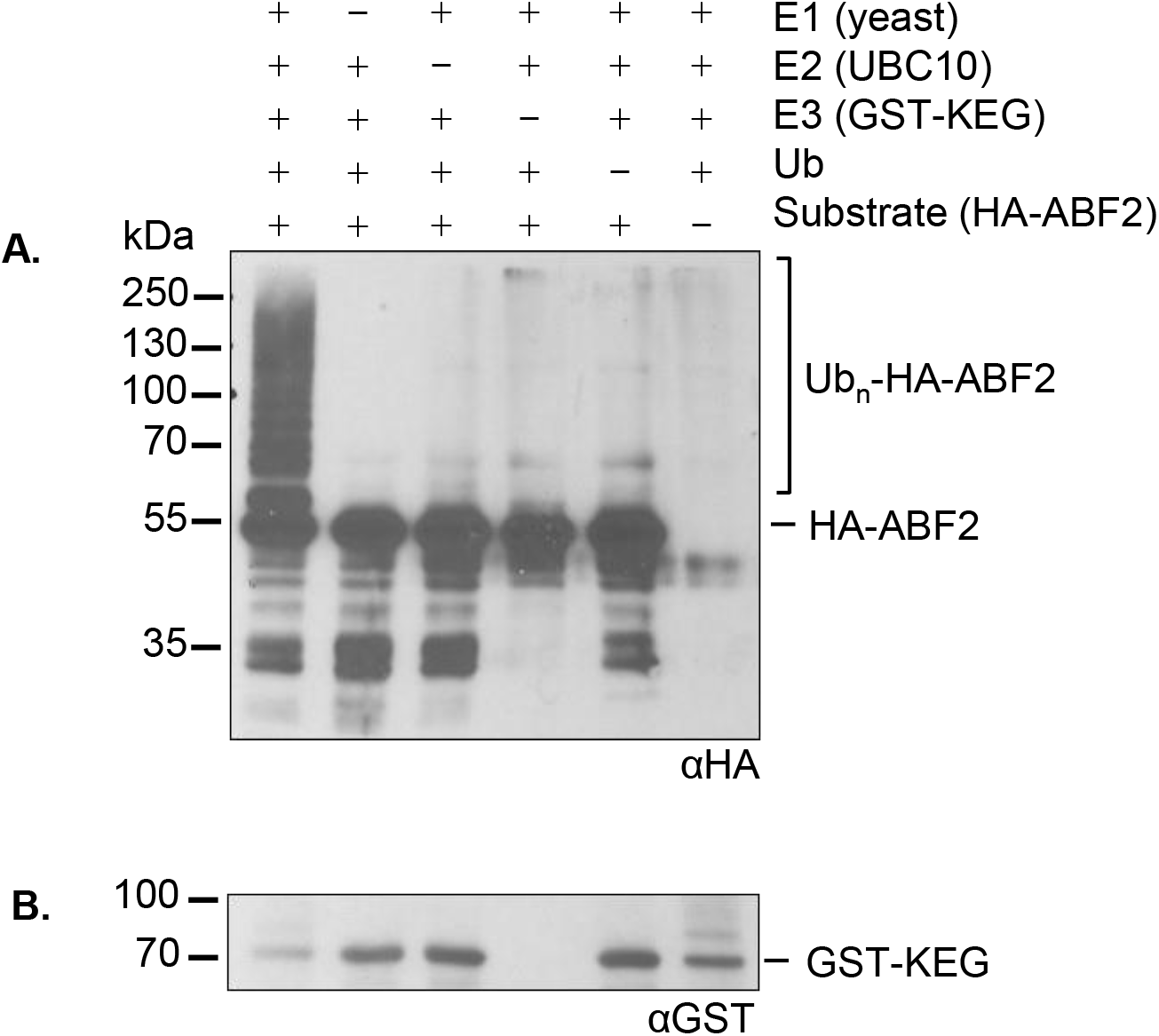
KEG-RKA ubiquitinates ABF2 *in vitro*. In an *in vitro* ubiquitination assay, recombinant His_6_-HA_3_-ABF2 was incubated with E1 (yeast recombinant), E2 (recombinant AtUBC10), the E3 GST-KEG-RK, and ubiquitin (Ub). After 1 hour, reactions were separated by SDS-PAGE. **(A)** His_6_-HA_3_-ABF2 was visualized with anti-HA western blotting. Components in the reaction are indicated with a +. **(B)** GST-KEG-RKA was visualized with anti-GST western blotting. Brackets indicate slower migrating forms of HA-ABF2 present only in the complete reaction, indicating ubiquitination.

To test whether KEG is important for ABF2 degradation, we used cell-free degradation assays to compare the degradation of bacterially expressed ABF2 protein in lysates from wild type seedlings to degradation in lysates from two different *keg* T-DNA alleles (*keg-1* and *keg-2*). *keg-1* contains a T-DNA insertion in the second exon and produces little or no *KEG* transcript, while *keg-2* contains a T-DNA insertion in the third intron and produces low levels of *KEG* transcript (Stone et al. 2006). Both lines exhibit similar seedling phenotypic differences from wild type (Stone et al. 2006). His_6_-HA_3_-ABF2 protein degraded in lysates from wild-type Col seedlings but was stable in lysates from both *keg-1* and *keg-2* seedlings (**Figures 4 and 5, respectively**) in parallel experiments with the same batch of recombinant protein. Less than 25% of His_6_-HA_3_-ABF2 protein remained after ten minutes in Col lysate, but there was no decrease in His_6_-HA_3_-ABF2 protein after ten minutes in either *keg* lysate.

**Figure 4:**
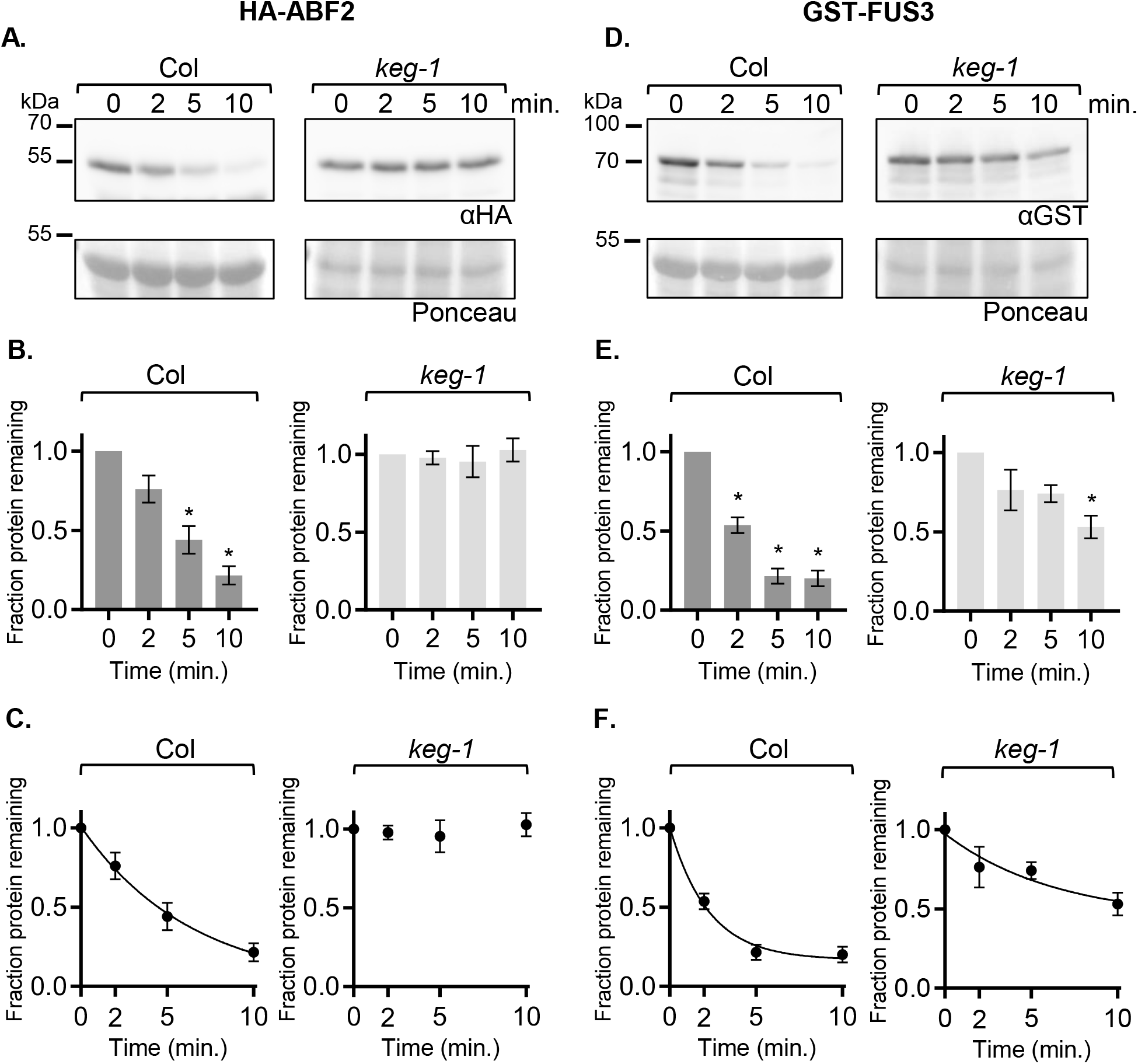
ABF2 protein is stabilized in *keg-1* lysate *in vitro*. In a cell-free degradation assay, recombinant His_6_-HA_3_-ABF2 or GST-FUS3 protein was added to lysate from Col or *keg-1* seedlings and the reaction was incubated at 22°C. Aliquots were removed at the indicated times, mixed with Laemmli sample buffer, and heated to 98°C to stop the reaction. **(A, D)** SDS-PAGE followed by anti-HA or anti-GST western blotting was used to visualize the His_6_-HA_3_-ABF2 or GST-FUS3 protein in each sample, respectively. Ponceau staining (below) was used to demonstrate equal loading. **(B, E)** Western blots were quantified with Image Studio Lite. Protein at time zero was normalized to 1 and the fraction of protein remaining at each timepoint was graphed. Asterisk (*) indicates the value differs significantly (p<0.05) from timepoint zero based on Anova (GraphPad Prism). **(C, F)** For each reaction, GraphPad Prism nonlinear regression was used to fit a curve modeling one-phase decay of His_6_-HA_3_-ABF2 or GST-FUS3. The *keg-1* line in **(C)** is not a one-phase decay curve because the data did not fit a decay curve. Bars are SE. N = 3 independent experiments (A and D show one representative experiment).

**Figure 5:**
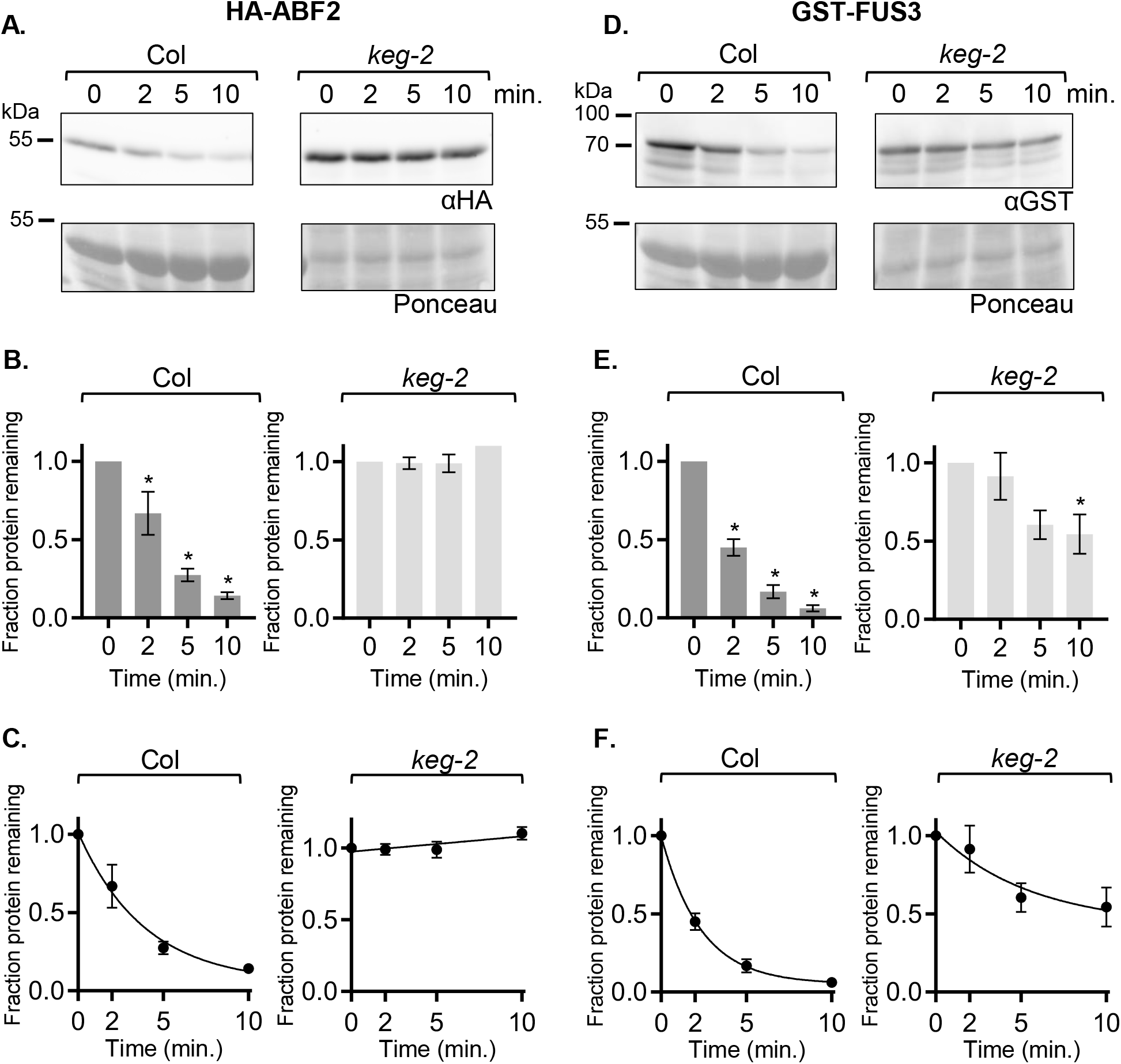
ABF2 protein is stabilized in *keg-2* lysate *in vitro*. In a cell-free degradation assay, recombinant His_6_-HA_3_-ABF2 or GST-FUS3 protein was added to lysate from Col or *keg-2* seedlings and the reaction was incubated at 22°C. Aliquots were removed at the indicated times, mixed with Laemmli sample buffer, and heated to 98°C to stop the reaction. **(A, D)** SDS-PAGE followed by anti-HA or anti-GST western blotting was used to visualize the His_6_-HA_3_-ABF2 or GST-FUS3 protein in each sample, respectively. Ponceau staining was used to demonstrate equal loading. **(B, E)** Western blots were quantified with Image Studio Lite. Protein at time zero was normalized to 1 and the fraction of protein remaining at each timepoint was graphed. Asterisk (*) indicates the value differs significantly (p<0.05) from timepoint zero based on Anova. (**C, F)** For each reaction, GraphPad Prism nonlinear regression was used to fit a curve modeling one-phase decay of His_6_-HA_3_-ABF2 or GST-FUS3. Bars are SE. N = 3 independent experiments (A and D show one representative experiment).

For the cell-free degradation assays described above, Col lysates were prepared from 7-day-old seedlings. To obtain developmentally comparable *keg* lysates, *keg* seedlings were grown for two weeks before harvesting, but were still smaller and paler than Col seedlings and rarely had true leaves. To demonstrate that the observed stability of ABF2 in *keg* lysates is not a property of the delayed development of *keg* seedlings and to further explore the substrate specificity of KEG, we tested whether *keg-1* lysates are capable of degrading other proteins. The cell-free degradation assay was used to compare the degradation of four additional transcription factors in both Col and *keg-1* lysates: the B3 transcription factor FUCSA3 (FUS3), the bHLH transcription factor ENHANCER OF GLABROUS 3 (EGL3), the bZIP transcription factor TGACG SEQUENCE-SPECIFIC BINDING PROTEIN 1 (TGA1, in bZIP group D) and the bZIP transcription factor G-BOX BINDING FACTOR 3 (GBF3, in bZIP group G) (**Figures 4-7**). FUS3 was selected because it is reported to be a proteasomal substrate that degrades rapidly in cell-free seedling degradation assays (Lu et al. 2010), and was previously used in our lab as a control for cell-free degradation assays with ABF1 and ABF3 (Chen et al. 2013). EGL3 is also reported to be a proteasomal substrate that degrades in seedling lysates (Patra et al. 2013). TGA1 and GBF3 were selected because they are bZIP transcription factors but are in different groups than the ABFs and affect different physiological processes, suggesting that their proteolytic regulation might be different (Dröge-Laser et al. 2018).

**Figure 6:**
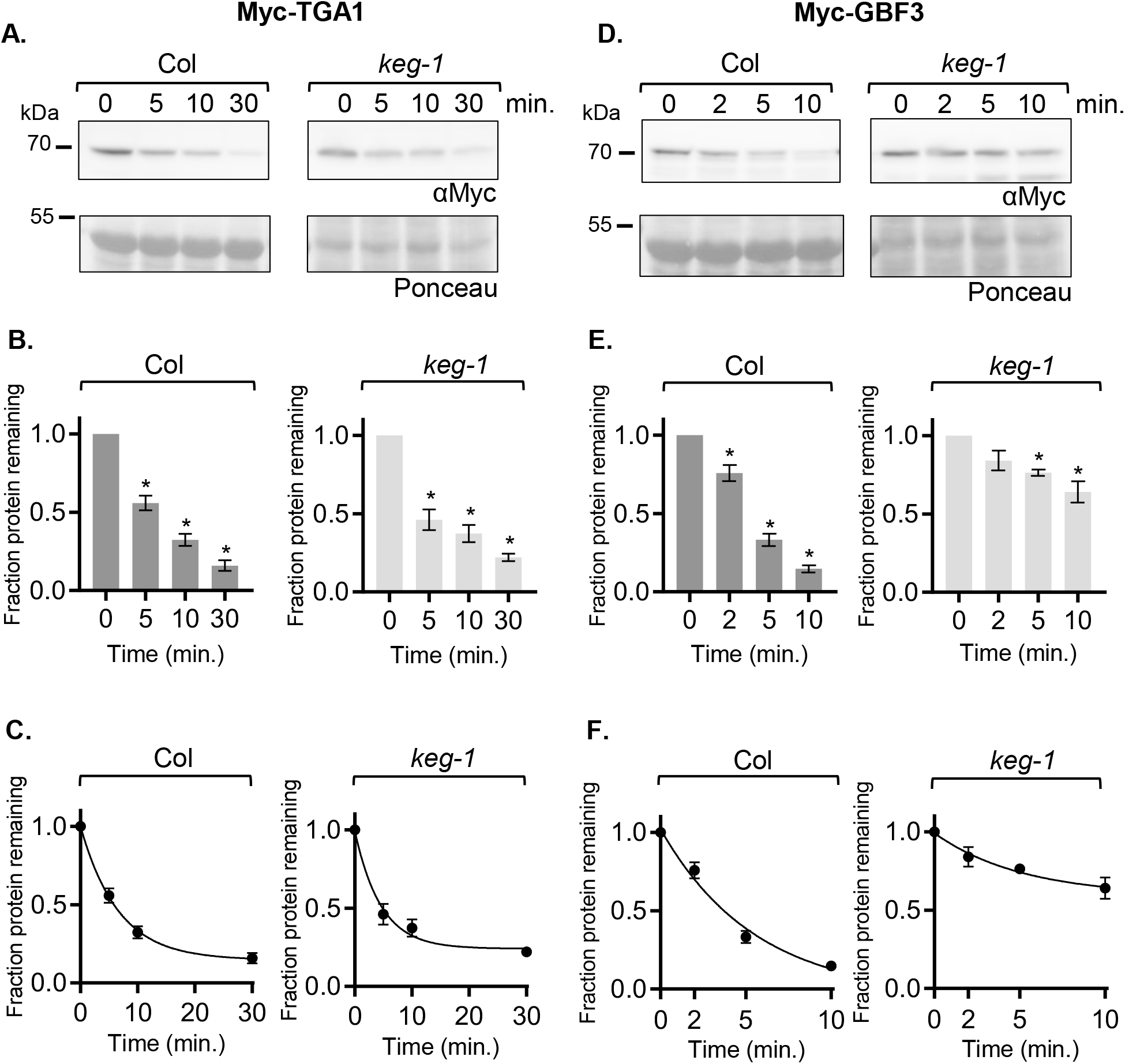
The bZIP proteins TGA1 and GBF3 degrade in *keg-1* lysate *in vitro*. In a cell-free degradation assay, recombinant Myc-TGA1 or Myc-GBF3 protein was added to lysate from Col or *keg-1* seedlings and the reaction was incubated at 22°C. Aliquots were removed at the indicated times, mixed with Laemmli sample buffer, and heated to 98°C to stop the reaction. **A, D)** SDS-PAGE followed by anti-Myc western blotting was used to visualize the Myc-TGA1 or Myc-GBF3 protein in each sample. Ponceau staining was used to demonstrate equal loading. **B, E** Western blots were quantified with Image Studio Lite. Protein at time zero was normalized to 1 and the fraction of protein remaining at each timepoint was graphed. Asterisk (*) indicates the value differs significantly (p<0.05) from timepoint zero based on Anova. **C, F)** For each reaction, GraphPad Prism nonlinear regression was used to fit a curve modeling one-phase decay of Myc-TGA1 or Myc-GBF3. Bars are SE. N = 3 independent experiments (A and D show one representative experiment).

**Figure 7:**
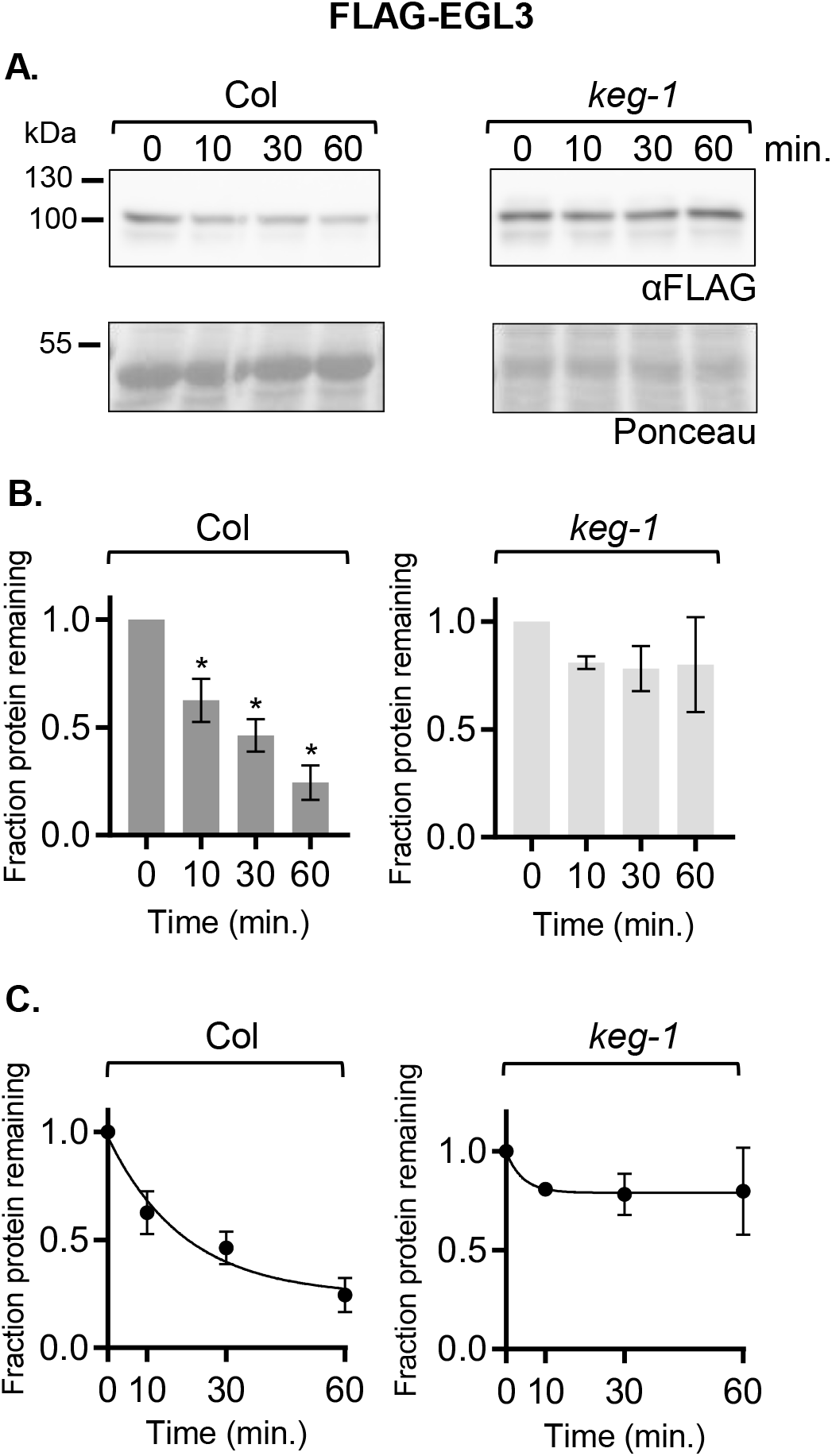
The bHLH protein EGL3 does not degrade significantly in *keg-1* lysate *in vitro*. In a cell-free degradation assay, recombinant FLAG-EGL3 protein was added to lysate from Col or *keg-1* seedlings and the reaction was incubated at 22°C. Aliquots were removed at the indicated times, mixed with Laemmli sample buffer, and heated to 98°C to stop the reaction. **(A)** SDS-PAGE followed by anti-FLAG western blotting was used to visualize the FLAG-EGL3 protein. Ponceau staining was used to demonstrate equal loading. **(B)** Western blots were quantified with Image Studio Lite. Protein at time zero was normalized to 1 and the fraction of protein remaining at each timepoint was graphed. Asterisk (*) indicates the value differs significantly (p<0.05) from timepoint zero based on Anova. **(C)** For each reaction, GraphPad Prism nonlinear regression was used to fit a curve modeling one-phase decay of FLAG-EGL3. Bars are SE. N = 3 independent experiments (A shows one representative experiment).

For all proteins with observed degradation, the data fit a pseudo-first order decay curve, as expected (graphs in C, **Figures 4-7**). TGA1 degraded at similar rates in Col and *keg-1* lysates (**Figure 6 A-C**). GBF3 and FUS3 also degraded in *keg-1* lysates, although their degradation was slower in *keg-1* lysates than in Col lysates (**Figures 4–5,D-F and 6 D-F, respectively**). Like ABF2, EGL3 was stabilized in *keg-1* lysates (**Figure 7**). Although the degradation of some proteins was slowed in *keg-1* lysates, these results show that protein degradation machinery is still functional in *keg-1* seedling lysates and that KEG is important for ABF2 degradation *in vitro*.

### *The C4 region affects ABF2 stability* in vitro

It was previously reported that a truncated ABI5 protein (ABI5^1-343^) lacking 99 C-terminal amino acids including the C4 region is more stable *in vitro* than full-length ABI5 (Liu and Stone 2013). To confirm that our cell-free degradation assay setup could replicate this result and to directly compare truncated ABFs to comparably truncated ABI5 proteins, we cloned and recombinantly expressed ABI5^1-343^ and ABI5-ΔC4, an ABI5 protein with a smaller deletion of just the C4 region (the 13 amino acids at the C-terminus). ABI5-ΔC4 is comparable to the ABF1 and ABF3 C4 deletions previously used in cell-free degradation assays (Chen et al. 2013). In cell-free degradation assays, His_6_-HA_3_-ABI5^1-343^ degraded more slowly than full-length His_6_-HA_3_-ABI5, as previously reported by Liu and Stone 2013 (**Figure 8**). There was significantly more ABI5^1-343^ protein than full-length ABI5 protein at the 2, 5, and 10 minute timepoints, and the predicted half-life of ABI5^1-343^ at 2.6 minutes was twice as long as the predicted half-life of full-length ABI5 at 1.2 minutes. His_6_-HA_3_-ABI5-ΔC4 also degraded more slowly than full-length His_6_-HA_3_-ABI5, but more quickly than His_6_-HA_3_-ABI5^1-343^ (**Figure 8**). The predicted half-life of His_6_-HA_3_-ABI5-ΔC4 was intermediate at 1.8 minutes and there was significantly more ABI5-ΔC4 protein than full-length ABI5 protein at the 5 and 10 minute timepoints.

**Figure 8:**
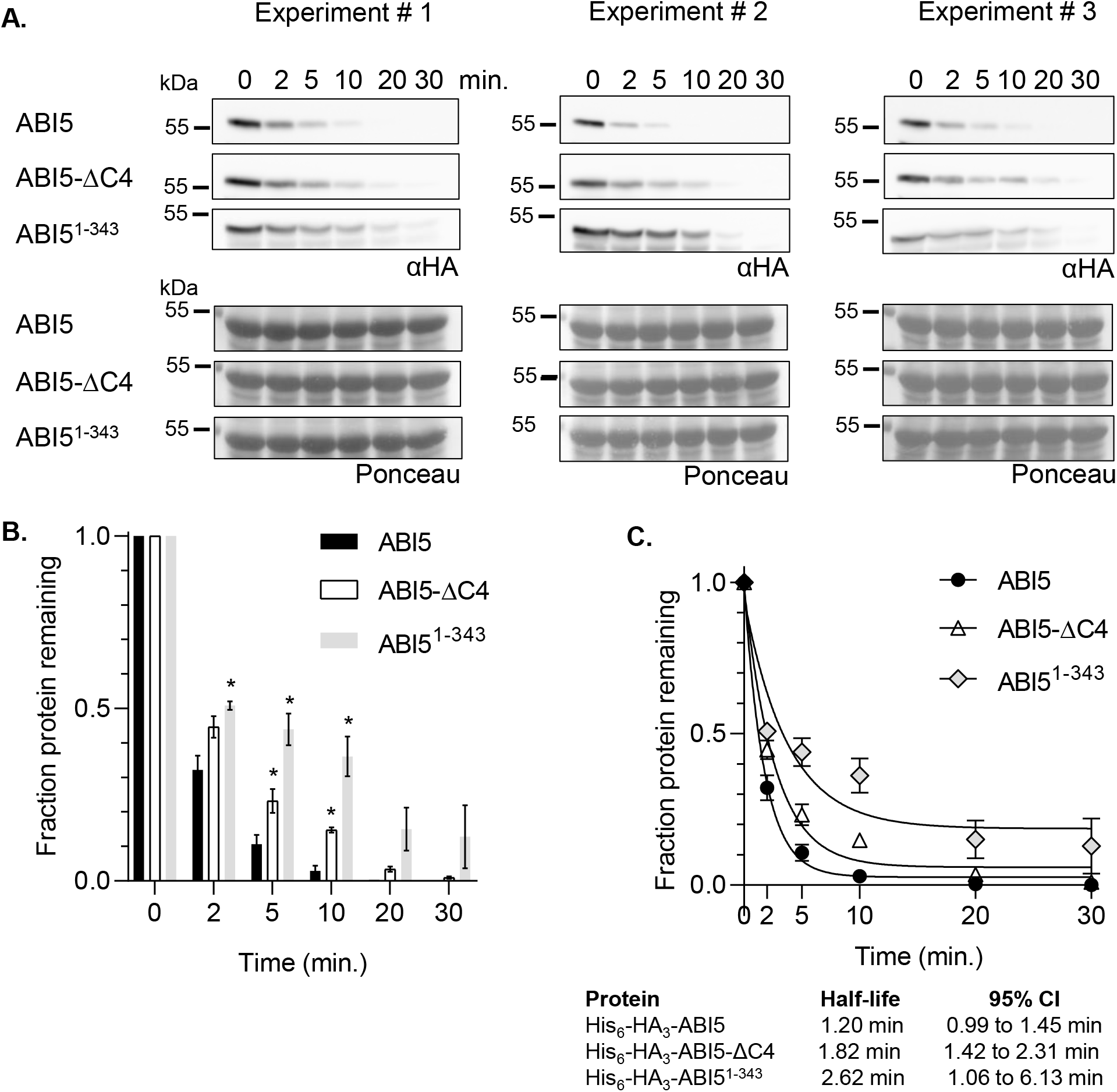
ABI5^1-343^ and ABI5-ΔC4 degrade more slowly than full-length ABI5 *in vitro*. In a cell-free degradation assay, recombinant His_6_-HA_3_-ABI5, -ABI5-ΔC4, or -ABI5^1-343^ protein was added to lysate from Col seedlings and the reactions were incubated at 22°C. Aliquots were removed at the indicated times, mixed with Laemmli sample buffer, and heated to 98°C to stop the reaction. **(A)** SDS-PAGE followed by anti-HA western blotting was used to visualize the HA-tagged protein in each sample. Ponceau staining (lower panels) was used to demonstrate equal loading. **(B)** Western blots were quantified with Image Studio Lite. Protein at time zero was normalized to 1 and the fraction of protein remaining at each timepoint was graphed. Asterisk (*) indicates significant difference (p<0.05) from fraction of full-length ABI5 protein remaining at the same timepoint based on Student’s t-test. Significance not tested for 20 and 30 minute timepoints. **(C)** For each reaction, GraphPad Prism nonlinear regression was used to fit curves modeling one-phase decay of each protein. R^2^ values for His_6_-HA_3_-ABI5, -ABI5-ΔC4, and -ABI5^1-343^ are 0.99, 0.98, and 0.86, respectively. Bars are SE. N = 3 independent experiments.

In contrast to ABI5, ABF1 and ABF3 proteins are less stable *in vitro* when their C4 regions are deleted (Chen et al. 2013) (see **Figure 9 A** for an alignment of the conserved regions in ABF proteins). To test whether ABF2 behaves similarly to ABI5 or ABF1 and ABF3, we cloned and recombinantly expressed a form of ABF2 lacking 12 amino acids at the C-terminus (ABF2-ΔC4). His_6_-HA_3_-ABF2-ΔC4 protein degraded more quickly than full length ABF2 in a cell-free degradation assay (**Figure 9**). The predicted half-life of full-length ABF2 at 3.8 minutes was longer than the predicted half-life of ABF2-ΔC4 at 2.1 minutes. Like ABF1 and ABF3, but in contrast to ABI5, ABF2 is stabilized by the C4 region.

**Figure 9:**
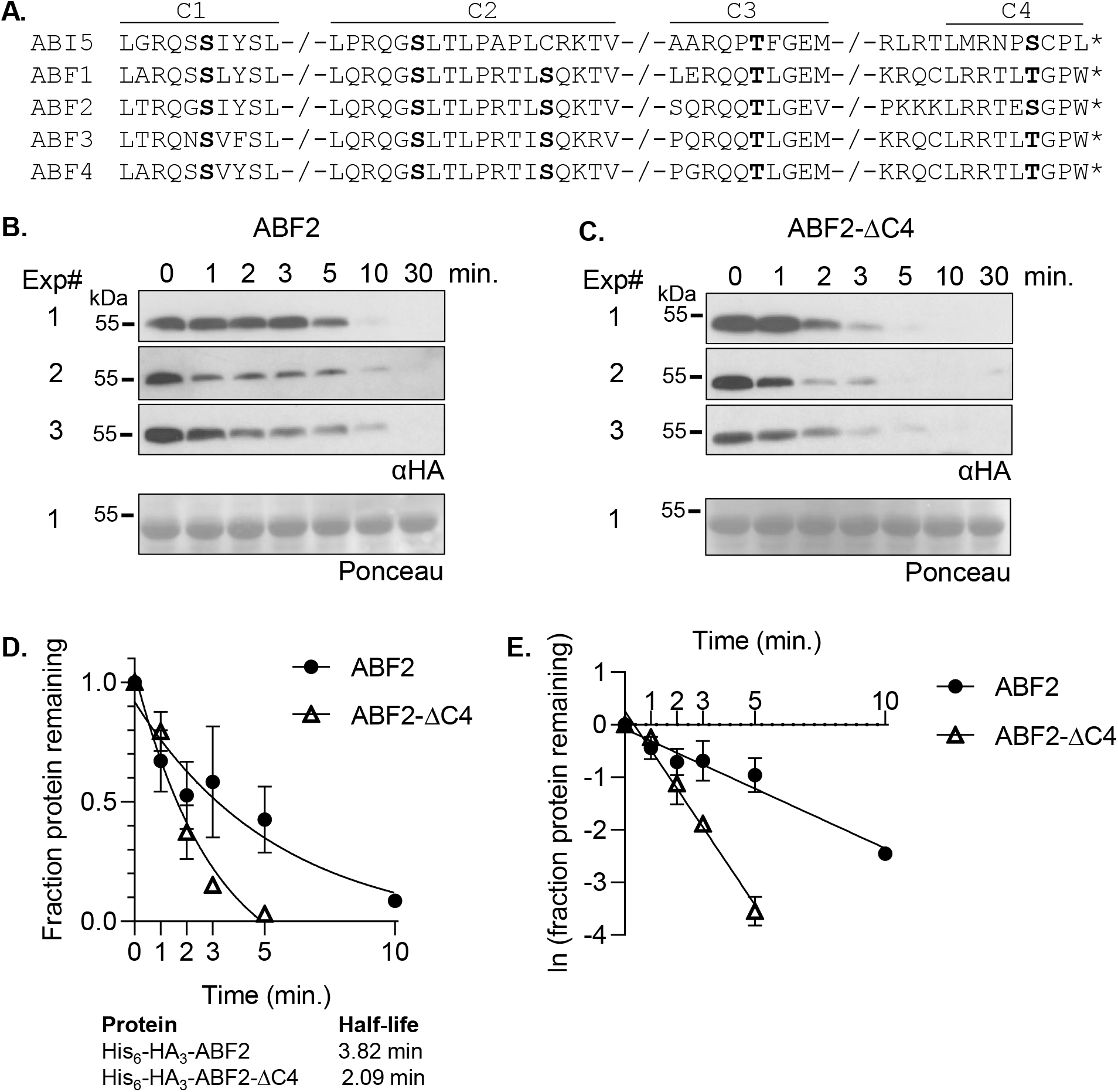
ABF2 lacking the C4 region degrades faster than full-length ABF2 *in vitro*. **(A)** Alignment of the conserved regions of ABF proteins. Phosphorylation sites are in bold. In a cell-free degradation assay, recombinant His_6_-HA_3_-ABF2 **(B)** or His_6_-HA_3_-ABF2-ΔC4 **(C)** was added to lysate from Col seedlings and the reaction was incubated at 22°C. Aliquots were removed at the indicated times, mixed with Laemmli sample buffer, and heated to 98°C to stop the reaction. Three independent experiments were performed. **(B, C)** SDS-PAGE followed by anti-HA western blotting was used to visualize the HA-tagged protein in each sample. Ponceau staining was used to demonstrate equal loading (Ponceau for Experiment #1 is shown). **(D)** Western blots were quantified with Image Studio Lite. Protein at time zero was normalized to 1 and the fraction of protein remaining at each timepoint was graphed. GraphPad Prism nonlinear regression was used to fit a curve modeling one-phase decay of each protein. Bars are SE. N = 3 independent experiments. **(E)** To compare slopes, a linear regression fit was performed in GraphPad Prism. Lines have significantly different slopes (p<0.01).

### Overexpression of full-length ABF1 or ABF1-ΔC4 causes delayed germination

Because overexpression of untagged ABF2 or ABF3 in Arabidopsis leads to delayed germination (Kang et al. 2002, Kim et al. 2004), we used this assay to determine whether the C4 region affects ABF function *in vivo*. We initially generated lines containing ABF2 FL and ABF2 delta C4 expressing transgenes, but rapid silencing in these lines prevented analyses. We were able to generate lines stably expressing versions of another ABF, ABF1, either full length (FL) His_6_-HA_3_-ABF1 or His_6_-HA_3_-ABF1-ΔC4 (**Figure 10A**) and evaluated the germination of independent homozygous transgenic lines over time compared to age-matched Col-0 control. We observed that seeds from all three FL ABF1 OE lines germinated more slowly than Col seeds on the same GM plates (**Figure 10 B**). After 32 hours, three lines overexpressing FL ABF1 reached 23.7%, 21.7%, and 0.6% germination, while Col (WT, age matched seeds) controls grown on the same plates reached 82.3%, 98%, and 94.8% germination, respectively. As a transformation control, germination of a transgenic line expressing GFP under the same *UBQ10* promoter in the same plasmid backbone was analyzed and its germination did not differ from that of untransformed Col (Supplemental Figure 5).

**Figure 10:**
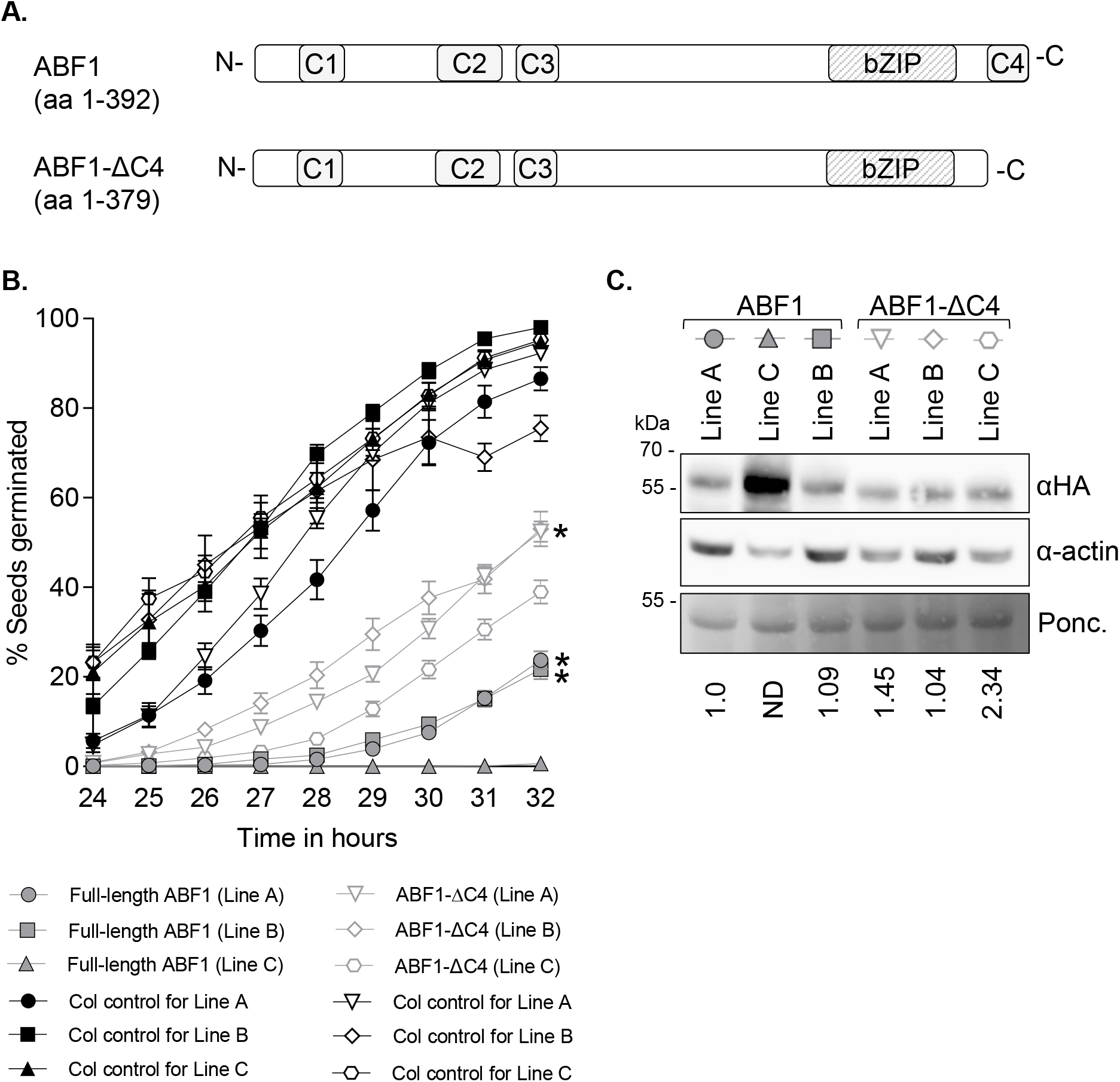
Overexpression of full-length ABF1 or ABF1-ΔC4 delays germination. **(A)** Diagram of full-length ABF1 (upper) and ABF1-ΔC4 (lower) proteins. **(B)** Seeds from plants overexpressing full-length ABF1 or ABF1-ΔC4 (or Col as a WT control on the same plate) were plated on GM, stratified at 4°C for 3 days, then incubated at 20°C under constant light. After plates were moved to 20°C, germination was scored hourly from 24 to 32 hours. Lines with similar protein expression levels are marked with an (*) (see panel C). N= at least 7 independent experiments, with 50 seeds per genotype per experiment. Bars are SE. **(C)** Comparison of HA-ABF1 protein level in each line. HA-tagged proteins were visualized with anti-HA western blotting (upper panel), and anti-actin western blotting (middle panel) was used as a loading control. Ponceau staining (lower panel) is an additional loading control. The same membrane was used for the anti-HA blot, anti-actin blot, and Ponceau stain. HA bands were normalized to actin bands, then compared to the lowest expressing full length line (ABF1 line A). ND = not determined (band not in linear range).

To test whether the C4 region affects ABF1’s ability to delay germination, we studied the germination of the ABF1-ΔC4 expressing lines. Germination of seeds from all three ABF1-ΔC4 lines was also slower than germination of the control Col seeds (**Figure 10 C, light gray lines and symbols**), with 52.4%, 53%, and 39% germination at 32 hours, compared to 92.3%, 75.5% and 95.3% for the Col controls. These results show that overexpression of the ABF1-ΔC4 protein affects germination. To more directly assess *in vivo* activity, the ABF1 protein levels between lines were directly compared using the HA tag (**Figure 10 C**). There seems to be a correlation between ABF1 protein and germination timing, with the higher expressing ABF1 FL line (Line C) exhibiting slower germination compared to ABF1 FL lines A and B. Similarly, among the ABF1-ΔC4 lines, the line with two-fold more expression (Line C) germinated more slowly than two other ABF1-ΔC4 lines (A and B). When comparing lines with equivalent expression of ABF1 FL vs ABF1-ΔC4, the ABF1-ΔC4 lines (Line A and Line B) germinated faster than the FL lines (Line A and Line B), suggesting that the ΔC4 version is less effective in slowing germination than the FL protein.

## DISCUSSION

ABF2 shares some aspects of proteolytic regulation with the other two previously analyzed ABF proteins in bZIP group A. These include instability both *in vitro* and in plants, accumulation regulated by the proteasome and ABA *in vivo*, *in vitro* ubiquitination by the E3 ligase KEG, KEG’s importance for their cell-free degradation, and the stabilizing effect of the C4 region *in vitro.* While most of these properties are consistent with proteolytic regulation of group A member ABI5, there are differences between the ABF proteins and ABI5. The C4 region destabilizes ABI5 *in vitro* but stabilizes ABF1, ABF2, and ABF3. ABF1-ΔC4, ABF2-ΔC4, and ABF3-ΔC4 proteins degrade faster than full-length proteins, while ABI5-ΔC4 degrades more slowly in parallel experiments.

Phosphorylation at a conserved site in the ABF3 C4 region (T451) is hypothesized to contribute to ABF3 stability by promoting ABF3 interaction with a 14-3-3 protein (Sirichandra et al. 2010). Whether S413 phosphorylation is important for other ABFs interactions with a 14-3-3 protein has not been published. The C-terminal amino acids of ABF1, 3, and 4 (LRRTLTGPW) and ABF2 (LRRTESGPW) contain both mode I (Rxx[S/T]xP) and mode II (Rxxx[S/T]xP) consensus 14-3-3 binding sites, while ABI5 (LMRNPSCPL), contains only a mode I site. However, in barley, HvABI5 interacts with several 14-3-3 proteins, and introducing a phospho-null substitution at T350 in HvABI5 eliminated interaction with 14-3-3 proteins in a yeast two-hybrid assay (Schoonheim et al. 2007), indicating that phosphorylation at the C4 site promotes 14-3-3 interaction with ABI5 as described for ABF3. Arabidopsis ABI5 also interacted with a 14-3-3 protein in a yeast two-hybrid assay (Jaspert et al. 2011). Because the C4 phosphorylation site appears to play similar role in 14-3-3 binding to ABI5 and ABF3, there might be other factors in the C4 region that account for the different role of the C4 in regulating ABI5 protein compared to the other ABFs.

There are other aspects of ABI5 proteolytic regulation that are likely to differ. ABI5 stability is regulated by S-nitrosylation, with modification of C-153 conferring enhanced degradation in a CUL4- and KEG-dependent manner (Albertos et al. 2015). ABF1-4 lack a corresponding cysteine residue at this position, suggesting that they are not regulated by NO in an analogous manner. Further studies are needed to dissect the differences and similarities between ABF1-4 and ABI5 proteolytic regulation.

In the cell-free degradation assay, we explored the ability of *keg* extracts to degrade other types of transcription factors and tested extracts from two different *keg* LOF alleles. Both *keg-1* and *keg-2* lysates exhibited identical properties. TGA1 degraded similarly in *keg-1* and Col lysates, indicating that *keg* lysates are capable of degrading proteins. We also observed that degradation of several non-ABF proteins was slower in *keg-1* lysates than in Col lysates. Degradation of GBF3 and FUS3 was slowed, and EGL3 was stable in *keg-1* lysates compared to Col lysates over the ten-minute time course. The FUS3 results were unexpected, because previous assays in our lab observed that FUS3 degraded equivalently in *keg-1* lysates (Chen et al. 2013). However, the timepoints used in Chen et al. 2013 extended from 30 to 90 minutes, and most FUS3 protein had degraded in both Col and *keg-1* lysate by the first timepoint at 30 minutes. Slower degradation at early timepoints might not have been observed in those experiments. Our FUS3 cell-free degradation assay extended from 1 to 10 minutes, at which point just over 53% of FUS3 protein remained.

*keg-1* seedlings have a very high level of ABI5 protein (Stone et al. 2006). The slowed degradation of FUS3 and GBF3 in *keg-1* lysate might result from altered ABA signaling in *keg-1,* since both FUS3 and GBF3 are involved in ABA responses and FUS3 degradation is ABA-dependent. GFP-tagged FUS3 protein increases following ABA treatment (Gazzarrini et al. 2004, Lu et al. 2010), and ABA inhibits FUS3 degradation *in vitro* (Chiu et al. 2016). Although the bZIP protein GBF3 is not in bZIP group A, it is also known to play a role in ABA response. Plants overexpressing GBF3 are more drought resistant than wild-type plants and exhibit less inhibition of root growth by ABA (Ramegowda et al. 2017). EGL3 is involved in trichome development and anthocyanin synthesis and its degradation is mediated by the HECT E3 ligase UBIQUITIN PROTEIN LIGASE 3 (UPL3) (Patra et al. 2013). It is unclear why EGL3 is stabilized in *keg-1* lysate.

The reduced stability of addtional proteins in *keg* extracts observed here is consistent with the pleotropic phenotype of *keg* mutants (Stone et al. 2006) and the observation that simultaneous removal of ABI5 and several ABF proteins in the *keg* background has only a minor effect on the *keg* phenotype (Chen et al. 2013). Identification of CIPK26 (Lyzenga et al. 2013), formate dehydrogenase (McNeilly et al. 2018), JAZ12 (Pauwels et al. 2015), and MKK4 and 5 (Gao et al. 2020) as KEG interactors/substrates further support the model that KEG has broad effects on plant growth and stress responses. The roles of KEG in immunity and vascular development implicate KEG in plant defense responses, intracellular trafficking (Gu and Innes 2011, Gu and Innes 2012), and cellular differentiation (Gandotra et al. 2013), in addition to its role in the regulation of group A bZIP proteins. These additional processes as well as downstream ABA responses affected in *keg* could indirectly modulate the proteolysis of other transcription factors.

## SUPPLEMENTAL MATERIALS

Supplemental Table 1 Primers for genotyping T-DNA lines

Supplemental Table 2 Primers for plant expression vector cloning

Supplemental Table 3 Primers for recombinant protein expression vector cloning

Supplemental Table 4 Transgenic line information

Supplemental Table 5 Primers for qPCR

Supplemental Figure 1 Additional replicas for CHX, MG132, and ABA experiments on ABF2 OE lines

Supplemental Figure 2. Additional constitutively expressed bZIP group A members accumulate in seedlings after incubation in MG132.

Supplemental Figure 3. Additional constitutively expressed bZIP group A proteins accumulate in seedlings after incubation in ABA.

Supplemental Figure 4. Constitutively expressed AREB3 bZIP group A protein accumulates in seedlings after incubation in ABA.

Supplemental Figure 5. Vector control for HA-ABF1 overexpression lines. Germination of seeds expressing GFP under control of the *UBQ10* promoter does not differ from germination of WT seeds.

## ACKNOWLEDGEMENTS

This work was supported by a grant from the National Science Foundation (IOS-1557760 to J. Callis, PI) and funds from the Paul K. and Ruth R. Stumpf Endowed Professorship in Plant Biochemistry and Aggie Alumni Foundation Award to J.C. We thank Eli Nambara and Ruth Finkelstein for the gift of *abi5-7* seeds and Dr. Finkelstein for assistance with genotyping the *abi5-7* allele and helpful discussions. We thank Sonia Gazarrini for the gift of the GST-FUS3 construct, Ling Yuan at the University of Kentucky for the *EGL3* in the pGEX-4T-1 vector, Sara Hotton for the synthesis of the modified pEXP1-DEST Myc9 expression vector (p7296), and Jonathan Gilkerson for synthesis of the modified pET3c expression plasmid (p3832). We thank the University of California-Davis Controlled Environment Facility (CEF) for assistance with the propagation of transgenic plants. We thank many past members of the Callis laboratory for helpful discussions, most recently John Riggs. We thank Jemina Cornejo, Eduards Norkvests, Anna Chen, and Stephanie Ng for assistance in propagation and analyses of transgenic lines.

## AUTHOR CONTRIBUTIONS

KL, Y-TC, and JC designed the research; KL, Y-TC, KK, and BS performed research; KL, Y-TC, JC, and BS analyzed data; KL and JC wrote the paper.

