## Supplemental Figures and Table for "Factors that affect protein abundance of the bZIP transcription factor ABRE-BINDING FACTOR 2 (ABF2), a positive regulator of abscisic acid signaling"

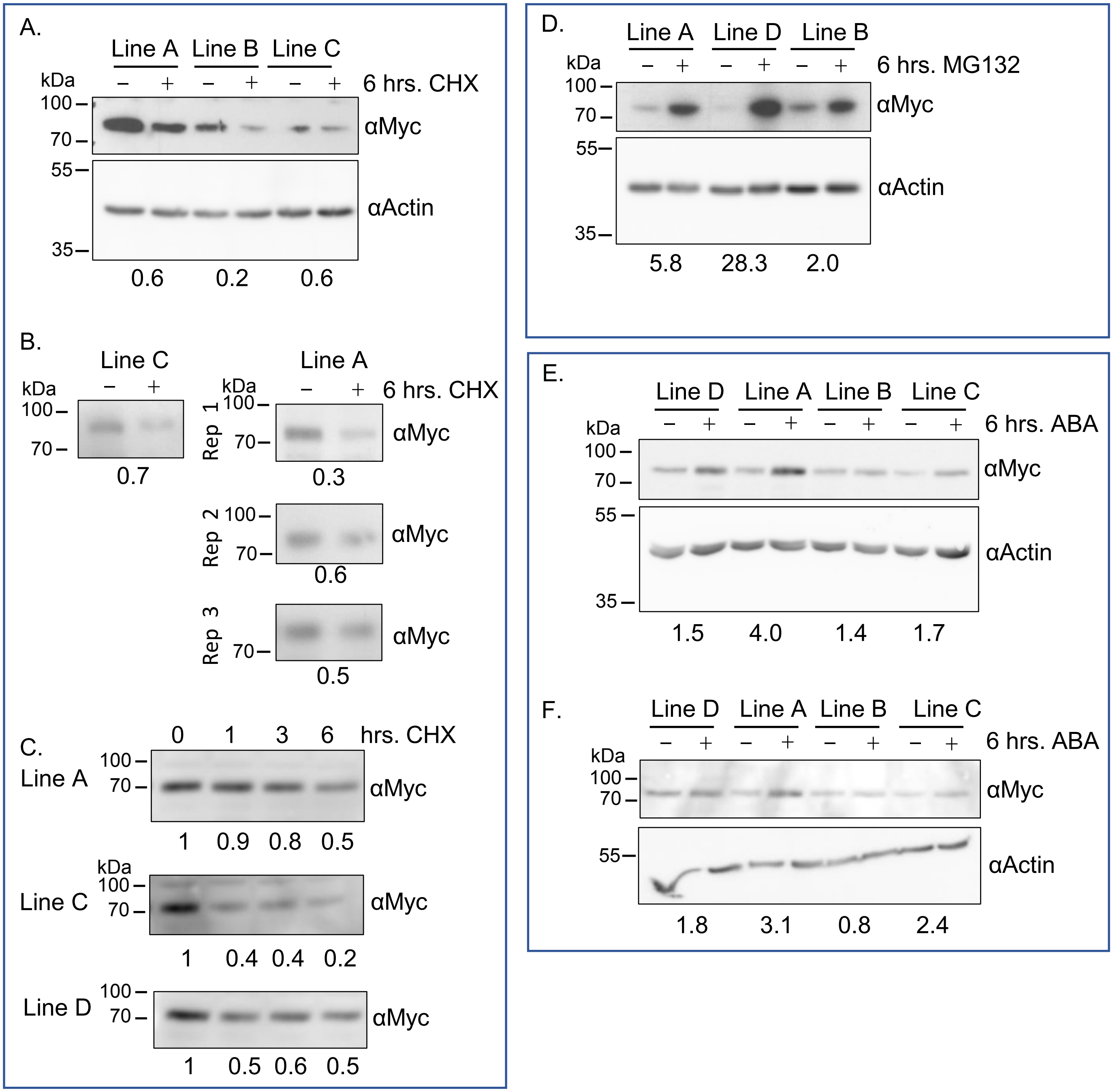


**Supplemental Figure 1. Additional replicas for CHX, MG132, and ABA experiments on ABF2 OE lines**

**A)** Additional independent replica of the CHX experiment shown in Figure 1A, with hmz T3 seedlings. **B)** Additional independent CHX experiments with hmz T4 seedlings, where Myc-ABF2 proteins were immunoprecipitated with anti-Myc beads. **C)** CHX time courses with T4 seedlings. **B,C)** No actin normalization. Values represent fraction remaining relative six hours of mock treatment, designated "0".

**D)** Additional independent replica of MG132 experiment shown in Figure 1B, with segregating T2 seedlings.

**E, F)** Additional independent replicas of the ABA experiment shown in Figure 1C, with T4 seedlings.

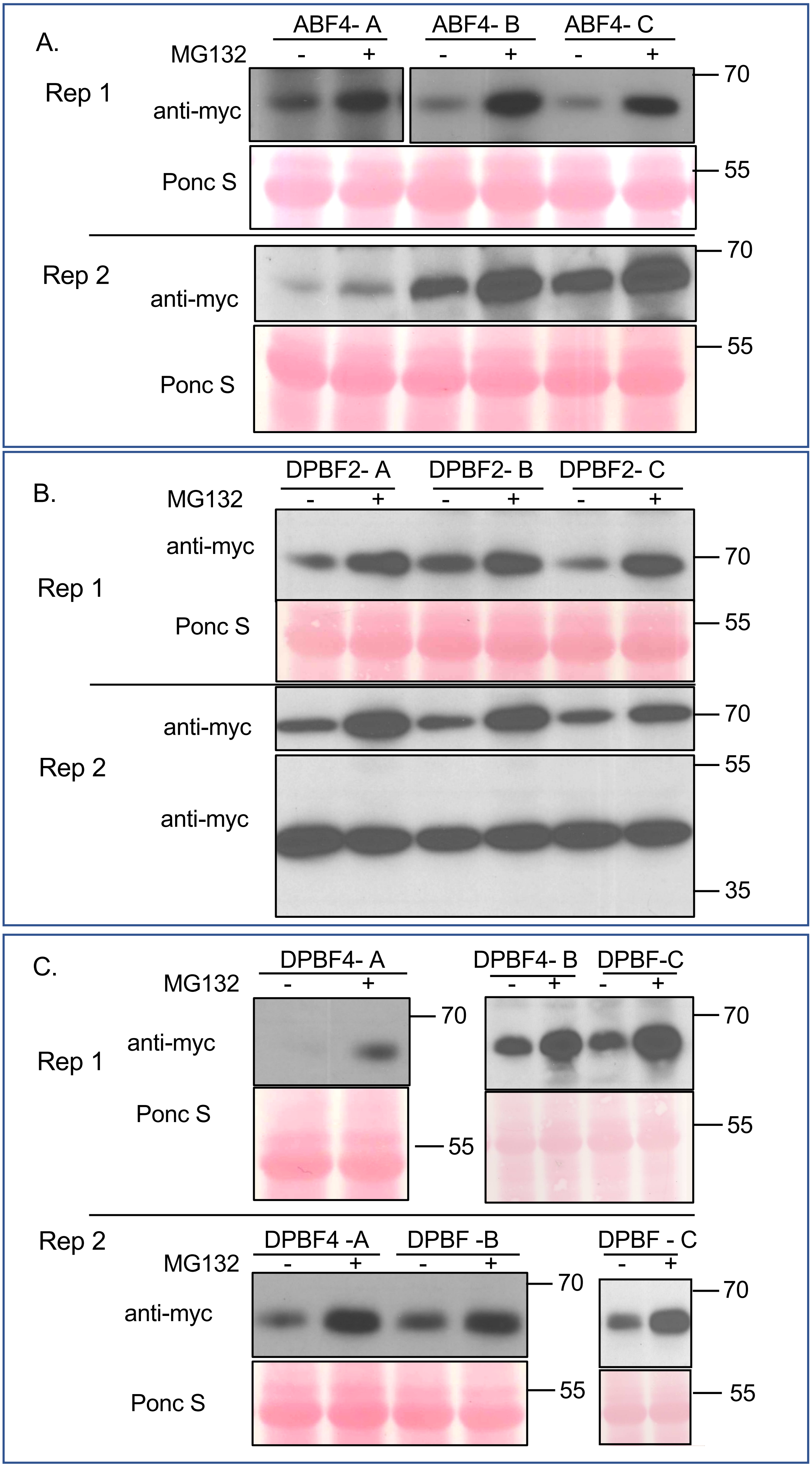
A.
ABF4- A
ABF4- B
ABF4- C
MG132
- + - + - +
Rep 1
anti-myc
Ponc S
70
55
Rep 2
anti-myc
Ponc S
70
55
B.
DPBF2- A
DPBF2- B
DPBF2- C
MG132
- + - + - +
Rep 1
anti-myc
Ponc S
70
55
Rep 2
anti-myc
anti-myc
70
55
35
C.
DPBF4- A
DPBF4- B
DPBF-C
MG132
- + - + - +
Rep 1
anti-myc
Ponc S
70
55
Rep 2
DPBF4 -A
DPBF -B
DPBF - C
MG132
- + - + - +
anti-myc
Ponc S
70
55

**Supplemental Figure 2. Additional constitutively expressed bZIP group A members accumulate in seedlings after incubation in MG132.**

7-day-old liquid grown T2 seedlings from three independent lines with 3:1 segregation for the transgene expressing 10xMyc-tagged ABF4 (A), DPBF2 (B) or DPBF4 (C) were treated with either 50 μM MG132 or 0.5% DMSO as a solvent control. After 6 hours, the seedlings were flash frozen in liquid nitrogen and ground in IP buffer. Equal amounts of total protein per sample (typically 50 μM total protein) were loaded for SDS-PAGE, and anti-Myc western blotting was used to visualize the Myc-tagged protein in each sample (upper panel). Two independent experiments (rep 1, rep 2) are shown, with the same line replicated shown directly underneath. To demonstrate equivalent loading, the membranes were stained with Ponceau S (lower panel). In (B) rep 2, a background anti-Myc band is shown as a loading control. In (A), immunoblot images represent the same gel, but two different exposure times were used; one for line A and a different time for lines B and C. For panel (C), breaks represent samples from the same experiment run on separate gels. Each rep indicates a separate growth, treatment and lysate.

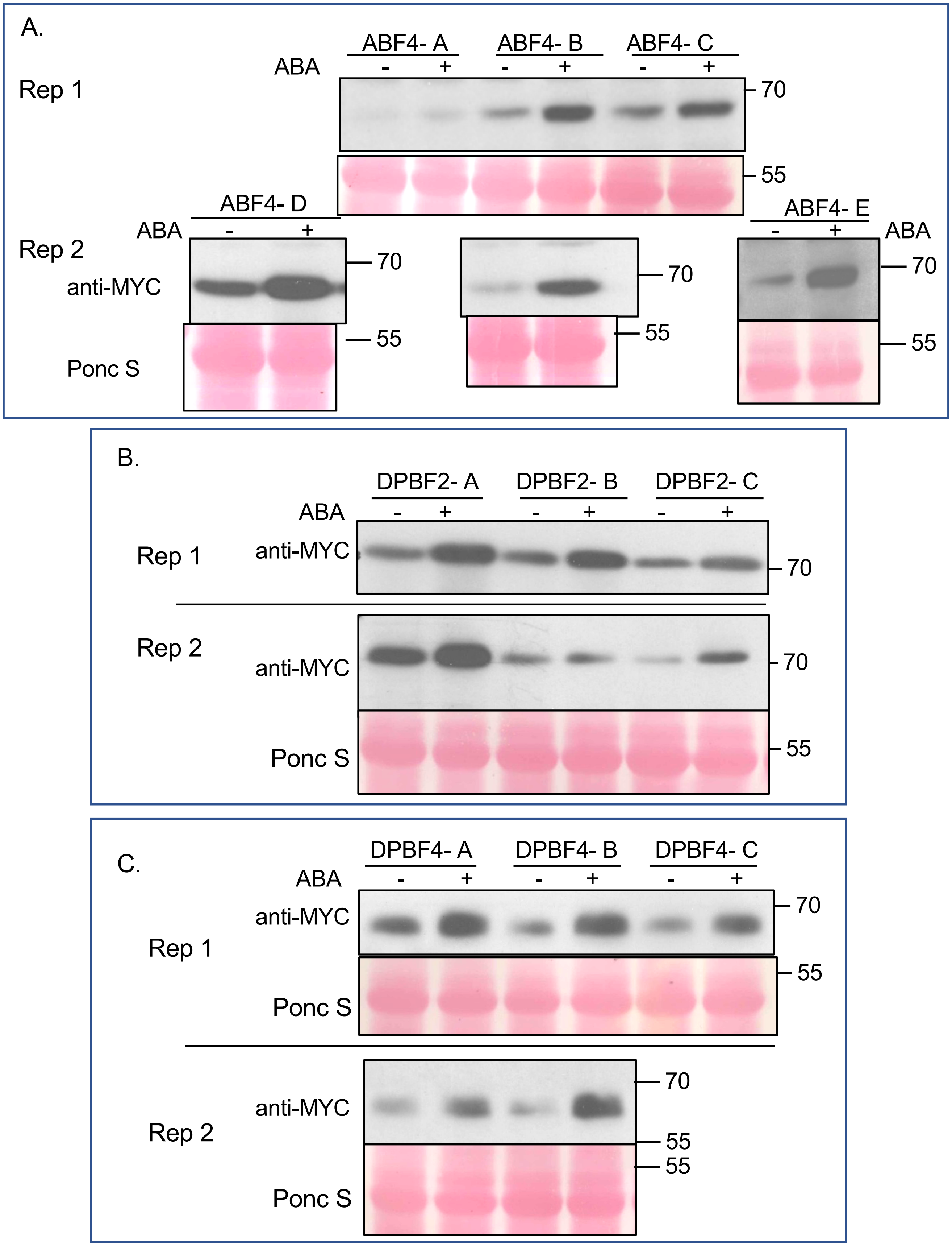
A.
ABF4- A
ABF4- B
ABF4- C
ABA
- + - + - +
Rep 1
70
55
ABF4- D
ABA
- +
Rep 2
anti-MYC
Ponc S
70
55
70
55
ABF4- E
- +
ABA
70
55
B.
DPBF2- A
DPBF2- B
DPBF2- C
ABA
- + - + - +
Rep 1
anti-MYC
70
Rep 2
anti-MYC
70
Ponc S
55
C.
DPBF4- A
DPBF4- B
DPBF4- C
ABA
- + - + - +
anti-MYC
70
Rep 1
Ponc S
55
anti-MYC
70
Rep 2
55
55
Ponc S

**Supplemental Figure 3. Additional constitutively expressed bZIP group A proteins accumulate in seedlings after incubation in ABA.**

7-day-old liquid grown T2 seedlings from at least three independent lines with 3:1 segregation for the transgene expressing 10xMyc-tagged ABF4 (A), DPBF2 (B) or DPBF4 (C) were treated with either 50 μM ABA dissolved in ethanol or with 0.5% ethanol as a solvent control. After 6 hours, the seedlings were flash frozen in liquid nitrogen and ground in IP buffer. Equal amounts of total protein per sample (typically 50 μM total protein) were loaded for SDS-PAGE, and anti-Myc western blotting was used to visualize the Myc-tagged protein in each sample (upper panel). Two independent experiments (rep 1, rep 2) are shown, with the same line replicated shown directly underneath. To demonstrate equivalent loading, the membrane was stained with Ponceau S (lower panel). In (A), rep 2, all samples were prepared at the same time: lines D and B were run on the same gel, but are re-arranged to show the same order as in rep 1. Line E was run on a separate gel. In (B) rep 1, there is no loading control. Each rep indicates a separate growth, treatment and lysate.

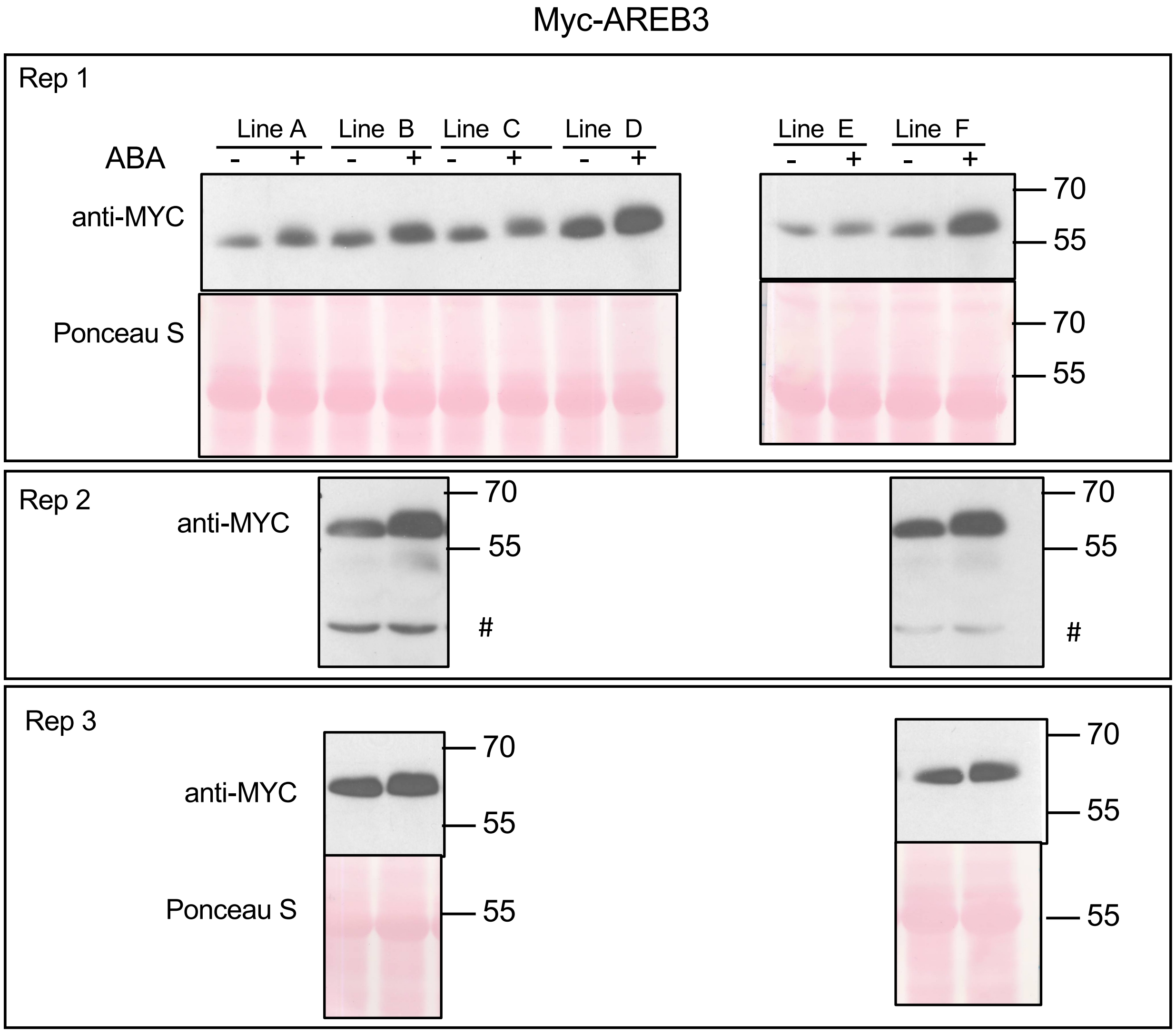


**Supplemental Figure 4. Constitutively expressed AREB3 bZIP group A protein accumulates in seedlings after incubation in ABA.**

7-day-old liquid grown T2 seedlings from at least three independent lines with 3:1 segregation for the transgene expressing 10xMyc-tagged AREB3 were treated with either 50 µM ABA dissolved in ethanol or with 0.5% ethanol as a solvent control. After 6 hours, the seedlings were flash frozen in liquid nitrogen and ground in IP buffer. Equal amounts of total protein per sample (typically 50 µM total protein) were loaded for SDS-PAGE, and anti-Myc western blotting was used to visualize the Myc-tagged protein in each sample (upper panel). Three independent experiments (rep 1-3) are shown, with the same line replicated shown directly underneath. To demonstrate equivalent loading, the membrane was stained with Ponceau S (lower panel). For rep 2, a non-specific band (#) is shown as a loading control. In rep 1, all samples were prepared at the same time and run on two separate gels. In reps 2 and 3 lines were run on the same gel, but an lane between was deleted. Each rep indicates a separate growth, treatment and lysate. Note that Line E (in rep 1) did not show an increase in ABA, this non-response was replicated in 2 additional experiments (not shown).

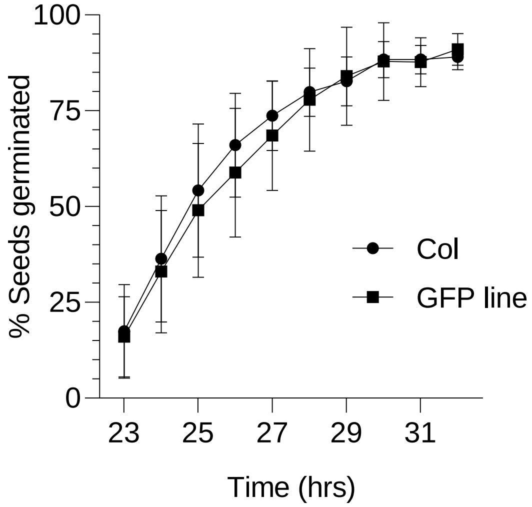


**Supplemental Figure 5. Vector control for HA-ABF1 overexpression lines. Germination of seeds expressing GFP under control of the *UBQ10* promoter does not differ from germination of WT seeds.**

Seeds from plants overexpressing GFP in the same vector as the HA-ABF lines in Figure 10 (or Col as a WT control on the same plate) were plated on GM, stratified at 4$^{o}$C for 3 days, then incubated at 20$^{o}$C under constant light. After plates were moved to 20$^{o}$C, germination was scored hourly from 24 to 32 hours. N= 12 independent experiments, with 50 seeds per experiment. Bars represent SD.

**Supplemental Table 1** Primers for genotyping T-DNA lines

| Callis Lab Primer # | AGI Locus Code | Gene Name | Primer Sequence (5' to 3') | Description |
|---|---|---|---|---|
| **5-253** | At5g13530 | *KEG* | TGTTCTTAAAAAGCTAGGCACGAA | Forward primer for genotyping SALK_049542 (*keg-1*). Use with 5-254. |
| **5-254** | At5g13530 | *KEG* | GGAGTGGATCCAGTGCACATC | Reverse primer for genotyping SALK_049542 (*keg-1*). Use with 5-253 for gene specific, and with 9-001 for T-DNA junction. |
| **9-113** | At5g13530 | *KEG* | GCAAGCGGCAATGCTGTGGTT | Forward primer for genotyping SALK_018105 (*keg-2*). Use with 9-114. |
| **9-114** | At5g13530 | *KEG* | AGAGCTTCCACCGCCTCCAGC | Reverse primer for genotyping SALK_018105 (*keg-2*). Use with 9-113 for gene specific, and with 9-001 for T-DNA junction. |
| **9-001** | - | - | TGGTTCACGTAGTGGGCCATCG | SALK T-DNA left border primer |

**Supplemental Table 2** Primers for plant expression vector cloning

| Callis Lab Primer # | AGI Locus Code | Gene Name | Primer Sequence (5' to 3') | Description |
|---|---|---|---|---|
| **9-363** | At3g19290 | *ABF4* | gggg aca agt ttg tac aaa aaa gca ggc tcg atgggaactcacatcaattt | 5' primer. ABF4-F. To amplify *ABF4* and clone into pDONR. |
| **9-364** | At3g19290 | *ABF4* | gggg ac cac ttt gta caa gaa agc tgg gtc tcaccatggtccggttaatg | 3' primer. ABF4-R. To amplify *ABF4* and clone into pDONR. |
| **9-365** | At3g56850 | *AREB3* | gggg aca agt ttg tac aaa aaa gca ggc tcg atggattctcagaggggta | 5' primer. AREB3-F. To amplify *AREB3* and clone into pDONR. |
| **6-366** | At3g56850 | *AREB3* | gggg ac cac ttt gta caa gaa agc tgg gtc tcagaaaggagccgagcttg | 3' primer. AREB3-R. To amplify *AREB3* and clone into pDONR. |
| **9-438** | At1g49720.1 | *ABF1* | gtca **attaat** atgggtactcacattgatat | 5' primer. ABF1-**AseI**-F, to amplify *ABF1* (both full-length and ΔC4) and clone into p3756. |
| **9-439** | At1g49720.1 | *ABF1* | gtca **ggatcc** ttaccacggaccggtaagg | 3' primer. ABF1-**BamHI**-R, to amplify full-length *ABF1* and clone into p3756. |
| **9-452** | At1g49720.1 | *ABF1* | gtca **ggatcc** ttaggccagcaatggaggctgctt | 3' primer. ABF1-ΔC4-**BamHI**-R, to amplify *ABF1-ΔC4* and clone into p3756. |

**Supplemental Table 3** Primers for recombinant protein expression vector cloning

| Callis Lab Primer # | AGI Locus Code | Gene Name | Primer Sequence (5' to 3') | Description |
|---|---|---|---|---|
| **6-1072** | At1g45249 | *ABF2* | TGAC**GGATCC**TCATGGAC<br>CTCCTTGCAGAAGATTCC | Reverse primer for cloning ABF2 deltaC4 (removing the C-terminal 13 amino acids), with 3' stop and **BamHI site**. |
| **6-1076** | At1g45249 | *ABF2* | **GGGGACCACTTTGTACAA**<br>**GAAAGCTGGGTC***GGATC*<br>*C* TCACCAAGGTCC | Reverse primer for cloning ABF2 into pDONR, with 3' *BamHI site* and **Gateway sequence**. |
| **6-1077** | At1g45249 | *ABF2* | **GGGGACCACTTTGTACAA**<br>**GAAAGCTGGGTC***GGATC*<br>*C*TCATGGACCTCC | Reverse primer for cloning His-HA-ABF2 deltaC4 into pDONR (same as 6-1076, but for deltaC4 which removes 39 bases from the C-terminus), with 3' *BamHI site*, stop, and **Gateway sequence**. |
| **6-1107** | At2g36270 | *ABI5* | CTGC **CATATG**<br>ATGGTAACTA<br>GAGAAACGAA G | 5' primer for cloning ABI5-FL, -$^{1-343}$, and -ΔC4, with 5' **NdeI site**. |
| **6-1108** | At2g36270 | *ABI5* | TGAT **GGATCC** TCA<br>CCTTCCCCTTAGCCCTCC | 3' primer for cloning ABI5$^{1-343}$ protein, with a stop after base 1029 and a 3' **BamHI site**. |
| **6-1109** | At2g36270 | *ABI5* | TGAT **GGATCC**<br>TTAGAGTGGACAACTC | 3' primer for cloning ABI5-FL, with a 3' **BamHI site**. |
| **6-1110** | At2g36270 | *ABI5* | TGAT **GGATCC** TCA<br>CCCGTTCGATTTCGGCAA<br>T | 3' primer for cloning ABI5-ΔC4, with a stop after base 1287 and a 3' **BamHI site**. |
| **9-440** | At1g45249 | *ABF2* | GTCA **CATATG**<br>GATGGTAGTATGAATTTG | ABF2-**NdeI**-F, forward primer for cloning *ABF2* into p3832. |
| **9-441** | At1g45249 | *ABF2* | gtca **ggatcc**<br>tcaccaaggtcccgactctgt | ABF2-**BamHI**-R, reverse primer for cloning *ABF2* into p3832. |
| **9-367** | At5g65210 | *TGA1* | gggg aca agt ttg tac aaa aaa<br>gca ggc tcg<br>atgaattcgacatcgacaca | TGA1-F. To amplify *TGA1* and clone into pDONR. |
| **9-368** | At5g65210 | *TGA1* | gggg ac cac ttt gta caa gaa<br>agc tgg gtc<br>ctacgttggttcacgatgtc | TGA1-R. To amplify *TGA1* and clone into pDONR. |
| **9-361** | At2g46270 | *GBF3* | gggg aca agt ttg tac aaa aaa<br>gca ggc tcg<br>atgggaaatagcagcgagg | GBF3-F. To amplify *GBF3* and clone into pDONR. |
| **9-362** | At2g46270 | *GBF3* | gggg ac cac ttt gta caa gaa<br>agc tgg gtc<br>tcagcctgcagctactgctt | GBF3-R. To amplify *GBF3* and clone into pDONR. |

**Supplemental Table 4** Transgenic line information

| Description | Gene Name | AGI Locus Code | Line name in manuscript | Callis Lab Name | Seedling Generation | Callis Lab Line # |
|---|---|---|---|---|---|---|
| Myc-ABF2 overexpression lines, with *35S* promoter | *ABF2* | At1g45249 | Line A | Line 2 | T2 (seg) | 12509-28 |
| | | | | | T3 (hmz) | 12702-2 |
| | | | | | T4 (hmz) | 13788 |
| | | | Line B | Line 5 | T2 (seg) | 12509-32 |
| | | | | | T3 (hmz) | 12704-4 |
| | | | | | T4 (hmz) | 13789 |
| | | | Line C | Line 1 | T2 (seg) | 12509-26 |
| | | | | | T3 (hmz) | 12705-3 |
| | | | | | T4 (hmz) | 13790 |
| | | | Line D | Line 3 | T2 (seg) | 12509-31 |
| | | | | | T3 (hmz) | 12897-4 |
| | | | | | T4 (hmz) | 13787 |
| $His_6$-$HA_3$-ABF1 overexpression lines, with *UBQ10* promoter | *ABF1* | At1g49720.1 | Line A | - | T4 (hmz) | 13152 |
| | | | Line B | - | T4 (hmz) | 13427 |
| | | | Line C | - | T4 (hmz) | 13975 |
| $His_6$-$HA_3$-ABF1-ΔC4 overexpression lines, with *UBQ10* promoter | *ABF1* | At1g49720.1 | Line A | - | T4 (hmz) | 13428 |
| | | | Line B | - | T4 (hmz) | 13431 |
| | | | Line C | - | T4 (hmz) | 13432 |
| Myc-ABF4 overexpression lines, with *35S* promoter | *ABF4* | At3g19290 | Line A | - | T2 (seg) | 12510-8 |
| | | | Line B | - | T2 (seg) | 12510-19 |
| | | | Line C | - | T2 (seg) | 12510-23 |
| | | | Line D | - | T2 (seg) | 12510-6 |
| | | | Line E | | T2 (seg) | 12510-16 |

**Supplemental Table 4 (continued)** Transgenic line information

| | | | | | | |
|---|---|---|---|---|---|---|
| Myc-DPBF2 overexpression lines, with *35S* promoter | *DPBF2* | At3g44460 | Line A | - | T2 (seg) | 12512-10 |
| | | | Line B | - | T2 (seg) | 12512-14 |
| | | | Line C | - | T2 (seg) | 12512-19 |
| Myc-DPBF4 overexpression lines, with *35S* promoter | *DPBF4* | At2g41070 | Line A | - | T2 (seg) | 12513-1 |
| | | | Line B | - | T2 (seg) | 12513-6 |
| | | | Line C | - | T2 (seg) | 12513-9 |
| Myc-AREB3 overexpression lines, with *35S* promoter | *AREB3* | At3g56850 | Line A | - | T2 (seg) | 12511-1 |
| | | | Line B | - | T2 (seg) | 12511-4 |
| | | | Line C | - | T2 (seg) | 12511-7 |
| | | | Line D | - | T2 (seg) | 12511-15 |
| | | | Line E | - | T2 (seg) | 12511-18 |
| | | | Line F | - | T2 (seg) | 12511-22 |
| GFP overexpression line, with *UBQ10* promoter | *GFP* | | GFP | - | T4 (hmz) | 13240 |

**Supplemental Table 5** Primers for qPCR

| Callis Lab Primer # | AGI Locus Code | Gene Name | Primer Sequence (5' to 3') | Description |
|---|---|---|---|---|
| **6-900** | - | - | GAACGGTAGCGCTGTTATCACAAG | 5' primer for *Myc-ABF2* qPCR. Located in the Myc tag, 5' of the *ABF2* sequence. |
| **7-755** | At1g45249 | *ABF2* | GAAACTCATCAAACGTCAACGA | 3' primer for *Myc-ABF2* qPCR. Starts 85 bases into the *ABF2* sequence. |
| **9-416** | At5g52310 | *RD29A* | CTTGATGGTCAACGGAAGGT | 5' primer for *RD29A* qPCR. Located at end of the 3rd exon. When used with 9-417, should span an 84-base intron. |
| **9-417** | At5g52310 | *RD29A* | CAATCTCCGGTACTCCTCCA | 3' primer for *RD29A* qPCR. Located at start of 4th exon. |
| **4-572** | At3g18780 | *ACT2* | AAGCTGGGGTTTTATGAATGG | 5' primer for *ACT2* qPCR. Located in the 3' UTR of *ACT2*, 24 bases after the stop codon. |
| **4-573** | At3g18780 | *ACT2* | TTGTCACACACAAGTGCATCAT | 3' primer for *ACT2* qPCR. Located in the 3' UTR of *ACT2*. |
